# GCK-4 regulates apical actin organization and lumen formation in the *C. elegans* intestine

**DOI:** 10.64898/2026.06.05.730387

**Authors:** Mohamed Attaibi, Jorian J. Sepers, João J. Ramalho, Ophélie Nicolle, Lynn Hasenöhrl, Savvas Tzavellas, Ruben Schmidt, Grégoire Michaux, Mike Boxem

**Author notes:** **Correspondence:** Mike Boxem.

## Abstract

Epithelial tubes are an essential component of many organ systems. The formation of their lumens depends on the close coordination of epithelial polarity, the apical actin cytoskeleton, and apical junctions, yet the mechanisms that organize the apical cytoskeleton and junctions downstream of polarity remain poorly understood. Here, we identify the Ste20 family kinase GCK-4, the single *C. elegans* ortholog of the mammalian kinases LOK and SLK, as a critical regulator of intestinal lumen formation. GCK-4 localizes to the apical membrane during early lumen formation and to microvillar tips as these structures develop. Its loss results in a lethal cystic lumen phenotype, without disrupting the establishment of epithelial polarity, and in occasional failure to maintain attachment between the pharynx and intestine. Lumens in *gck-4* mutant animals show irregular junction patterning, severely impaired apical accumulation of both actin and the membrane–actin linker Ezrin/Radixin/Moesin protein ERM-1, and microvilli atrophy. In *Drosophila* and mammalian cells, the GCK-4 orthologs are thought to organize the apical actin network largely through phosphorylation of ERM proteins. In contrast, ERM-1 phosphorylation is only partially reduced in *gck-4* mutants and is not abolished, indicating that GCK-4 acts through additional targets and is not the principal ERM-1 kinase in the intestine. Together, these findings identify GCK-4 as a key regulator of apical actin and junction organization during lumen formation, and demonstrate that Ste20 kinases can control apical cytoskeletal architecture at least partially independently of ERM phosphorylation.

## Introduction

Many organs are composed of epithelial tubes that transport liquids and gases. Epithelial lumen systems range in complexity from single-cell tubes, like the *C. elegans* excretory canal, to complex branched structures like mammalian lungs or mammary glands (Lubarsky and Krasnow, 2003; Sigurbjörnsdóttir et al., 2014). One of the mechanisms through which an epithelial tube can form is termed cord-hollowing. In this process, a lumen is created between apposing cells in a cord. This involves the formation of small cavities at the lumen initiation site which then expand and coalesce into a single continuous lumen (Datta et al., 2011; Lubarsky and Krasnow, 2003; Sigurbjörnsdóttir et al., 2014).

Lumen formation through cord-hollowing depends on the close coordination of the apical–basal polarity machinery, the apical actin cytoskeleton, and apical junctions. Formation of an apical domain determines where the lumen will form, and lumen initiation and growth depend on the delivery of proteins and lipids by the polarized trafficking machinery (Blasky et al., 2015; Buckley and St Johnston, 2022; Datta et al., 2011; Jewett and Prekeris, 2018). The actin cytoskeleton supports polarized trafficking and forms a dynamic cortical scaffold that generates the protrusive and tension-balancing forces needed to shape the lumen (Bovyn and Haas, 2024; Chan and Hiiragi, 2020; Navis and Bagnat, 2015). Apical junctions, in turn, distribute and modulate these forces between neighboring cells, prevent the paracellular passage of molecules, and help maintain segregation between apical and basolateral compartments (Buckley and St Johnston, 2022; Clarke and Martin, 2021; Matter and Balda, 2014; Mukenhirn et al., 2024; Shin et al., 2006). While the establishment of epithelial polarity is comparatively well understood, the mechanisms that organize the apical cytoskeleton and junctions during lumen formation remain far less defined.

The *C. elegans* intestine provides an excellent model to study the mechanisms that drive multicellular tube formation (Asan et al., 2016; Dimov and Maduro, 2019; Leung et al., 1999; Shaye and Soto, 2021; Zhang et al., 2017). The intestine is an epithelial tube consisting of 20 intestinal “E” cells that derive from a single E blastomere and are arranged in 9 rings (Int 1–9). The anterior ring, which is attached to the pharynx, consists of 4 cells, and the following rings consist of 2 opposing cells each. Formation of the intestine is a complex morphological process involving oriented divisions, cell intercalations, and circumferential rotation of int rings 2-4 (Asan et al., 2016; Leung et al., 1999), which take place as the intestine grows and elongates in concert with the whole embryo.

Lumen formation in the *C. elegans* intestine begins with the polarization of the intestinal primordium at the 8-cell stage (E8), when the apical polarity regulator PAR-3 and the junctional protein HMR-1 (E-cadherin) form punctate local polarity complexes (LPCs) at homotypic cell–cell contacts (Feldman and Priess, 2012; Naturale et al., 2023). As LPCs mature, they recruit additional apical components before coalescing at the midline by the late E16-stage, thereby defining the lumen initiation site. The PAR proteins PAR-3, PAR-6, and aPKC, the small GTPase CDC-42, and the adhesion protein HMR-1 are all required to establish or maintain a continuous apical domain at the midline (Pickett et al., 2022; Sallee et al., 2021). Following establishment of the lumen initiation site, physical lumen opening initiates at multiple sites along the midline, visible as separation between junctions of opposing cells, which expand and coalesce to form a single continuous lumen by the comma stage of embryonic development (Asan et al., 2016; Leung et al., 1999).

The molecular mechanisms that organize the apical cytoskeleton and junctions downstream of polarity, and drive the opening and shaping of the lumen, are far less well understood. A central effector is the membrane–cytoskeleton linker ERM-1, the single *C. elegans* ortholog of the mammalian ezrin, radixin, and moesin (ERM) proteins (Göbel et al., 2004; Van Fürden et al., 2004). ERM-1 is thought to couple the microvillar actin core to the plasma membrane, and its loss produces a cystic intestinal lumen together with reduced microvillar length and density (Göbel et al., 2004; Van Fürden et al., 2004). A second layer of regulation is provided by the subapical intermediate filament network: loss of the intermediate filament regulators *ifo-1*, *sma-5*, or *bbln-1* causes the apical domain to invaginate into the cytoplasm of intestinal cells (Carberry et al., 2009; Carberry et al., 2012; Coch and Leube, 2016; Geisler et al., 2020; Geisler et al., 2023). These defects, however, emerge during larval development, indicating a role in maintaining lumen integrity rather than in initial lumen formation. How the apical junctions themselves are patterned and repositioned as the lumen opens, and how this is coordinated with cytoskeletal organization, remains comparatively unexplored.

Here, we identify the Ste20 family kinase GCK-4, the single *C. elegans* ortholog of the mammalian kinases LOK (lymphocyte-oriented kinase) and SLK (Ste20-like kinase), as an essential regulator of intestinal lumen formation. GCK-4 localizes to the apical midline of the developing intestine and later to the tips of microvilli, and its loss produces a lethal cystic lumen with irregular junction patterning, severely impaired apical actin and ERM-1 accumulation, and microvillus atrophy. In *Drosophila* and mammalian cells, the GCK-4 orthologs LOK and SLK/Slik are thought to organize the apical actin network largely through phosphorylation of ERM-family proteins. However, ERM-1 phosphorylation is only partially reduced in animals lacking GCK-4, indicating that GCK-4 organizes apical actin through additional targets and is not the principal ERM-1 kinase in the intestine. Our findings establish GCK-4 as a key organizer of the apical actin cytoskeleton and cell junctions during lumen formation, and show that Ste20 kinases can control apical architecture independently of ERM phosphorylation.

## Results

### GCK-4 is required for intestinal development and localizes to the apical domain

To uncover novel regulators of intestinal lumen formation, we conducted a small-scale RNAi feeding screen targeting 10 kinases with putative roles in the intestine based on expression pattern or previously described functions of homologs in other species (Table 1). For each kinase, we performed RNAi by feeding on a strain expressing fluorescently tagged ERM-1 to visualize the morphology of the intestinal lumen. Feeding RNAi for *gck-4* resulted in lumen morphology defects, as well as in growth retardation and larval arrest. We therefore selected *gck-4* for further investigation. The *gck-4* gene encodes a member of the germinal center kinase-V (GCK-V) subfamily of the sterile 20p-like (Ste20) family of serine/threonine kinases, and is orthologous to the mammalian kinases LOK and SLK (Figure 1A and S1A). To investigate the function of *gck-4*, we constructed a *gck-4* deletion mutant lacking the complete coding region, referred to here as *gck-4(null)* (Figure S1B). Over 98% of *gck-4(null)* mutants arrest in early larval development (Figure 1B), and the intestines of *gck-4(null)* mutant larvae are cystic and show multiple lumen constrictions (Figure 1C, D). The presence of constrictions suggests that the L1 arrest may be due to an inability to take up sufficient nutrients. Consistent with this, feeding of fluorescently labelled dextran showed that the intestinal lumen of *gck-4(null)* mutants is not continuous (Figure S1C). Nevertheless, a very small percentage of animals survive to eventually become fertile adults, making it possible—with difficulty—to maintain *gck-4(null)* mutants as a homozygous strain.

**Figure 1:**
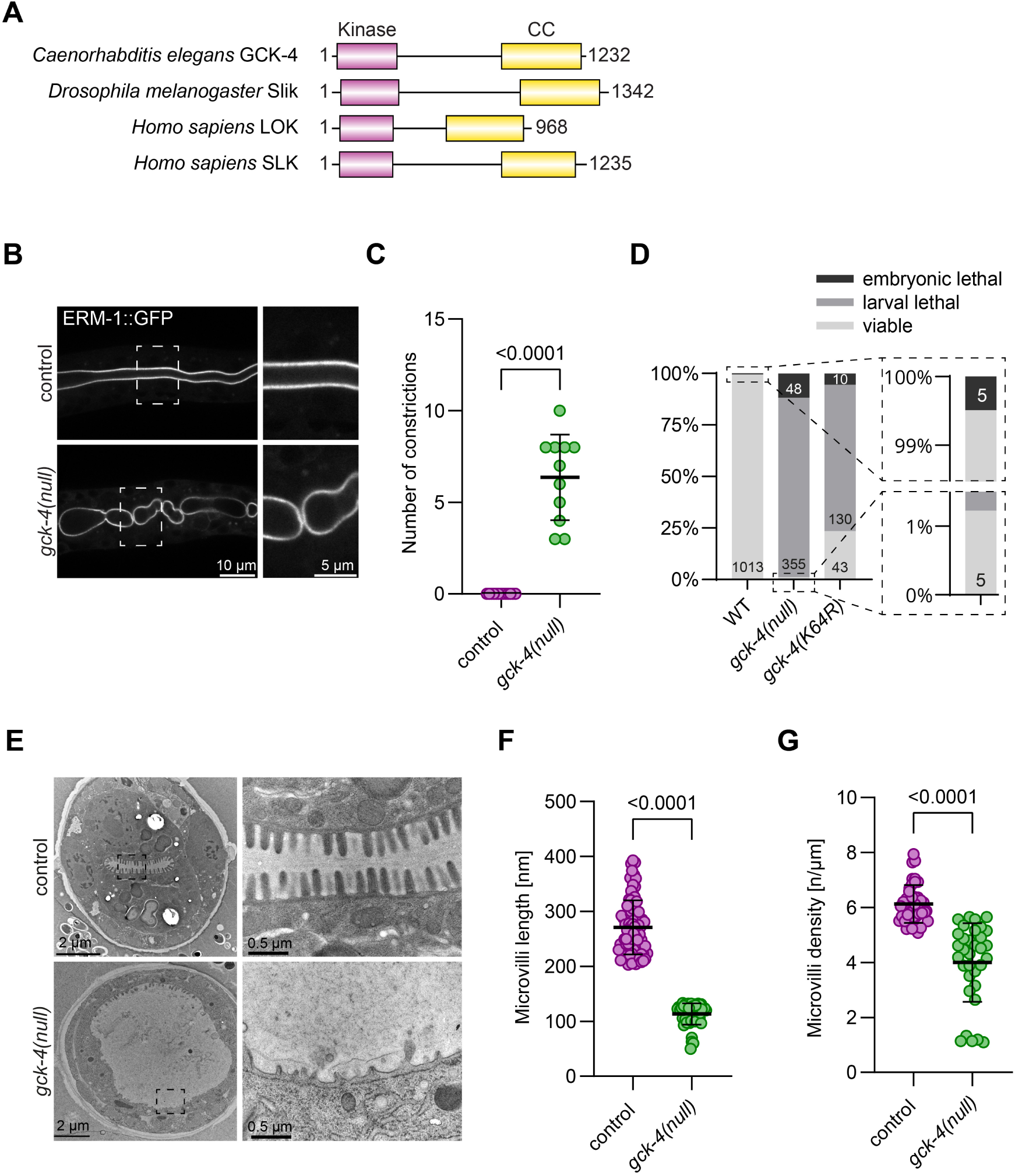
GCK-4 regulates intestinal lumen formation in ***C. elegans***. (A) Protein domain structure of GCK-4 and its orthologs in *Drosophila* and human. Protein length is in amino acids. (B) Lumen morphology in *gck-4(null)* mutant animals compared to control. ERM-1::GFP marks the microvillar brush border. Images show a single confocal microscopy plane. Enlarged versions of the boxed regions are shown to the right of each panel. Strains used: BOX1030 and BOX1065. (C) Number of lumen constrictions in L1 larvae. A constriction was counted when no separation was observed at the midline in YFP::ACT-5 signal. Each data point is the number of constrictions counted in a single worm. Data are represented as mean ± SD and analyzed with a Mann-Whitney test. Strains used: BOX1031 and BOX1059. (D) Embryonic lethality, larval arrest and survival to adulthood observed in progeny of animals of indicated genotypes. Data are represented as a proportion of the total amount of progeny, with the total number per outcome indicated in the graph. Total number of progeny counted in order of samples in graph: 1018, 408 and 183. Strains used: CGC1, BOX1036, BOX1402. (E) Transmission Electron Microscope (TEM) images of newly hatched *C. elegans* L1 larvae. Enlarged versions of the boxed regions are shown to the right of each panel. Strains used: N2 and BOX1036. (F-G) Quantifications of microvilli length and density from TEM images. Data are represented as mean ± SD and analyzed with Mann-Whitney test; p value shown on graph. Total number of animals analyzed in order of samples in graphs: 5 and 3 animals. For the 5 control animals, 42 measurements were obtained for microvilli density and 84 measurements for microvilli length. For the 3 *gck-4(null)* mutants, 33 measurements were obtained for microvilli density and 53 measurements for microvilli length.

**Table 1:**
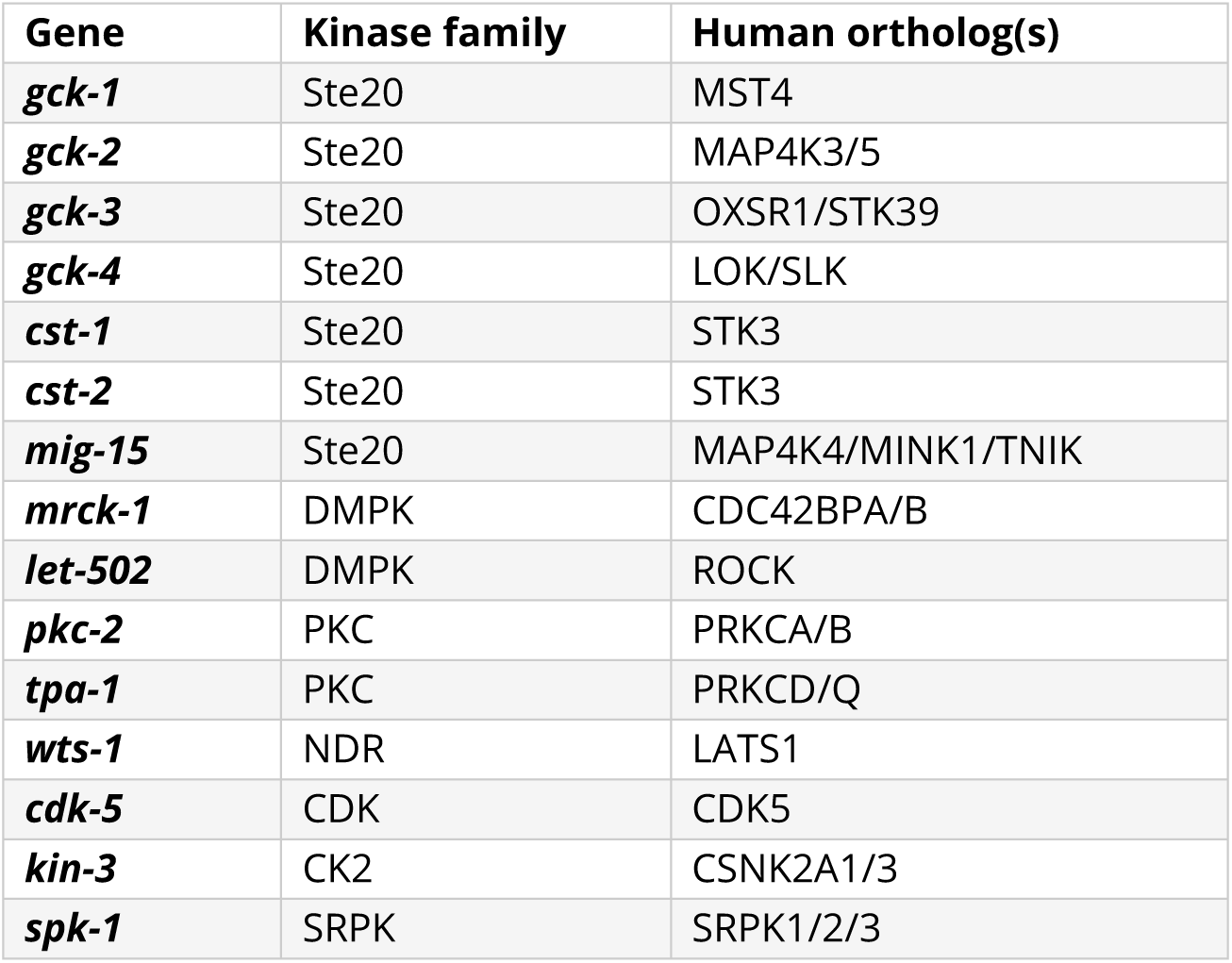
RNAi feeding screen targeting intestinal kinases.

The GCK-4 protein consists of a C-terminal kinase domain and an N-terminal coiled-coil domain separated by an unstructured central domain (Figure 1A). Members of the GCK-V subfamily of Ste20 kinases in other systems are involved in a range of biological processes, and some of their functions appear to be independent of catalytic activity (Pelaseyed et al., 2017; Rambaud et al., 2025). To investigate if the intestinal phenotypes in *gck-4(null)* mutants result from the loss of kinase activity, we constructed a kinase-dead *gck-4(K64R)* mutant by changing the lysine at position 64 to an arginine (K64R) in the endogenous locus. We also constructed a GFP-tagged (K64R) variant, which still localizes apically, indicating that the K64R mutation does not affect GCK-4 protein localization (Figure S2A, B). Similar to the *gck-4(null)* mutant, we observed cystic intestines and larval lethality in *gck-4(K64R)* animals (Figure 1D, S1E). Nevertheless, *gck-4(K64R)* mutants were more viable than *gck-4(null)* mutants. Thus, while much of the activity of GCK-4 is mediated through phosphorylation of downstream targets, GCK-4 may have additional kinase-independent functions.

To investigate the luminal defects of *gck-4* mutants in more detail, we performed transmission electron microscopy imaging on L1-stage *gck-4(null)* mutant animals. Wild-type L1 control larvae showed the typical oval lumen shape with a densely packed microvillar array, as well as electron dense apical junctions at cell-cell interfaces and intermediate filament endotube surrounding the lumen (Figure 1E). As expected, *gck-4(null)* lumens were irregular in shape and enlarged. In addition, the brush border of *gck-4(null)* animals displayed far fewer and shorter microvilli than controls (Figure 1F, G). Moreover, we observed defects in the appearance of apical junctions, including longer junctions, and irregular clustered distribution of the electron dense material (Figure S1D). The intermediate filament endotube, however, appeared unaffected. In summary, ultrastructural imaging shows that GCK-4 is important not only for the overall size and shape of the lumen but also for the formation of the brush border and cell junctions.

To determine the expression and subcellular localization patterns of GCK-4, we next constructed an endogenously expressed GCK-4::GFP fusion protein. GCK-4::GFP was expressed in multiple epithelial tissues, including the intestine, pharynx, excretory canal, the seam cells, spermatheca, and vulva (Figure 2 and S2C). In the intestine, GCK-4::GFP was expressed from early bean stage onwards, localizing to the apical domain (Figure 2A). To determine the precise subcellular location of GCK-4 in the intestine during larval stages, we compared the localization of GCK-4::GFP to ERM-1::mCherry, which localizes along microvilli (Ramalho et al., 2020). GCK-4 localized apical to ERM-1, demonstrating that GCK-4 is restricted to the microvillar tip region (Figure 2B, C).

**Figure 2:**
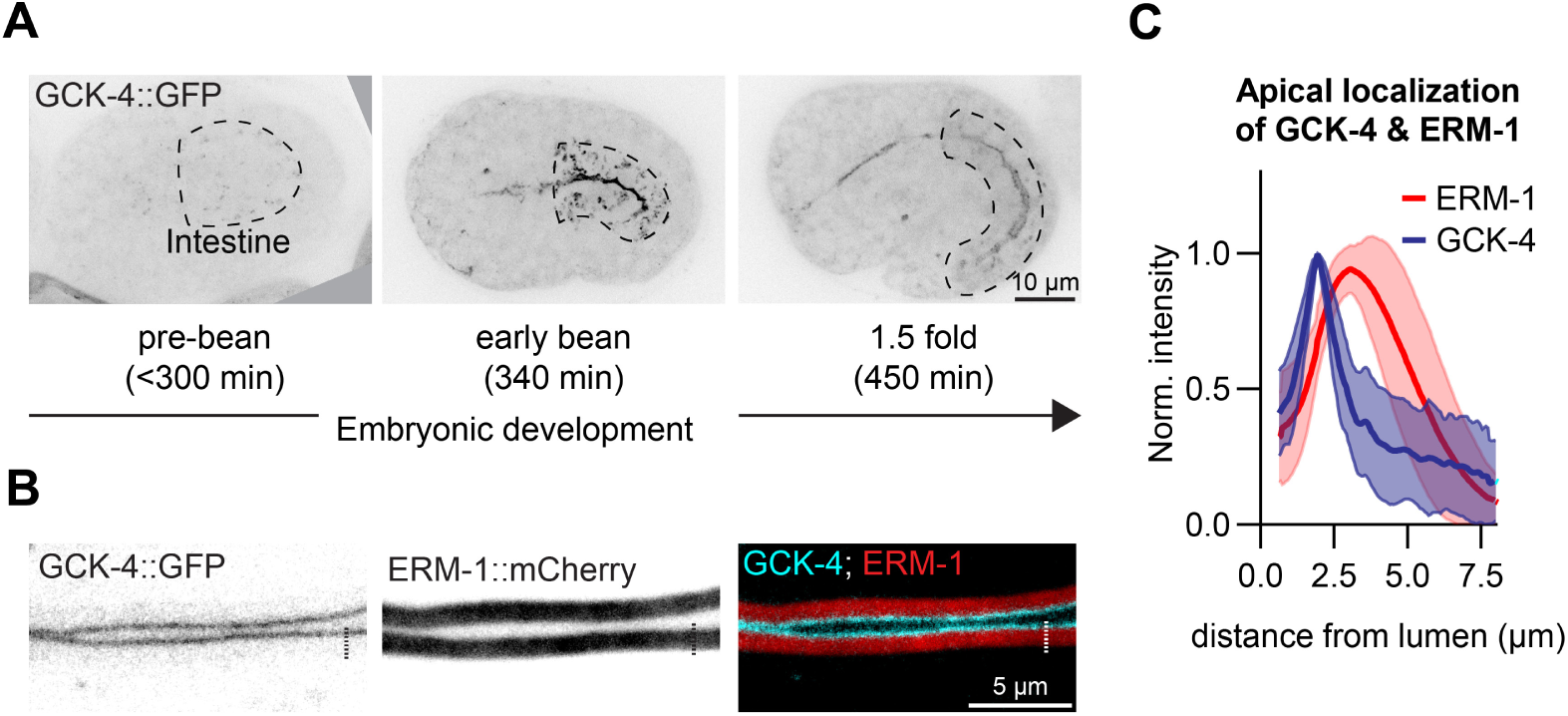
GCK-4 localizes to the apical luminal membrane domain during embryonic development and to the microvilli tips in larvae. (A) Maximum intensity projections of 3D stacks taken with a spinning disc confocal microscope showing the localization of GCK-4::GFP in embryonic development. Dashed lines outline the intestine. Time indicates approximate time in minutes after fertilization at 20–22°C according to WormAtlas (Hall and Altun, 2009). Strain used: BOX909. (B) Single focal plane taken with a spinning disc confocal microscope showing localization of GCK-4::GFP and ERM-1::mCherry::AID* in L4 larvae. Dashed line indicates example of a line used to quantify apical localization of GCK-4 and ERM-1 in (C). Strain used: BOX1095. (C) Quantification of apical localization of GCK-4 and ERM-1. Samples have been normalized to peak in signal and aligned to peak in GCK-4 signal. Data are represented as mean (line) ± SD (shaded area). *N* = 13. Strain used: BOX1095.

The expression of GCK-4 in the excretory canal prompted us to examine if GCK-4 loss also causes defects in this tubular organ. Indeed, excretory canal length was decreased in freshly hatched *gck-4(null)* larvae than in controls, indicating a general role for GCK-4 in lumen formation (Figure S2D–E).

In summary, our results establish GCK-4 as an essential regulator of intestine formation that localizes to the luminal domain during early intestinal development and to the tips of microvilli in larval stages.

### GCK-4 is not required for the establishment of intestinal epithelial polarity

One of the earliest steps in the formation of the *C. elegans* intestinal tract is the establishment of apical-basal polarity in the intestinal primordium. Similar to GCK-4 loss, intestine-specific depletion of the apical polarity regulators PAR-3, PAR-6, aPKC, and CDC-42 results in the hatching of larvae with cystic obstructed intestines leading to early larval arrest (Pickett et al., 2022; Sallee et al., 2021). We therefore determined whether *gck-4* mutants are defective in intestinal epithelial polarity establishment.

As early as the E8 stage of intestinal development, the apical polarity regulator PAR-3 and the junctional protein HMR-1 (E-cadherin) break cellular symmetry by forming local polarity complexes (LPCs) at homotypic cell–cell contacts. As LPCs mature they recruit additional components, including the apical proteins PAR-6 and PKC-3 (aPKC) and the junctional protein AFD-1, before coalescing at the midline of the E16-stage intestinal primordium (Naturale et al., 2023). To determine if GCK-4 is involved in these early steps of intestinal polarization, we followed the localization of PAR-6::GFP and HMR-1::GFP in *gck-4(null)* mutant animals. In both control and *gck-4(null)* animals, PAR-6 and HMR-1 were first apparent in foci that presumably are LPCs, and subsequently localized to the midline of the gut (Figure 3A, B). By the E16 stage, PAR-6::GFP localized along the entire intestinal midline in *gck-4(null)* mutant embryos (Figure 3C). Together, these results indicate that both symmetry breaking and the initial establishment of a luminal domain at the central midline can occur independently of *gck-4*. Thus, we conclude that the intestinal defects found in *gck-4* mutants are not due to defects in early polarity establishment.

**Figure 3:**
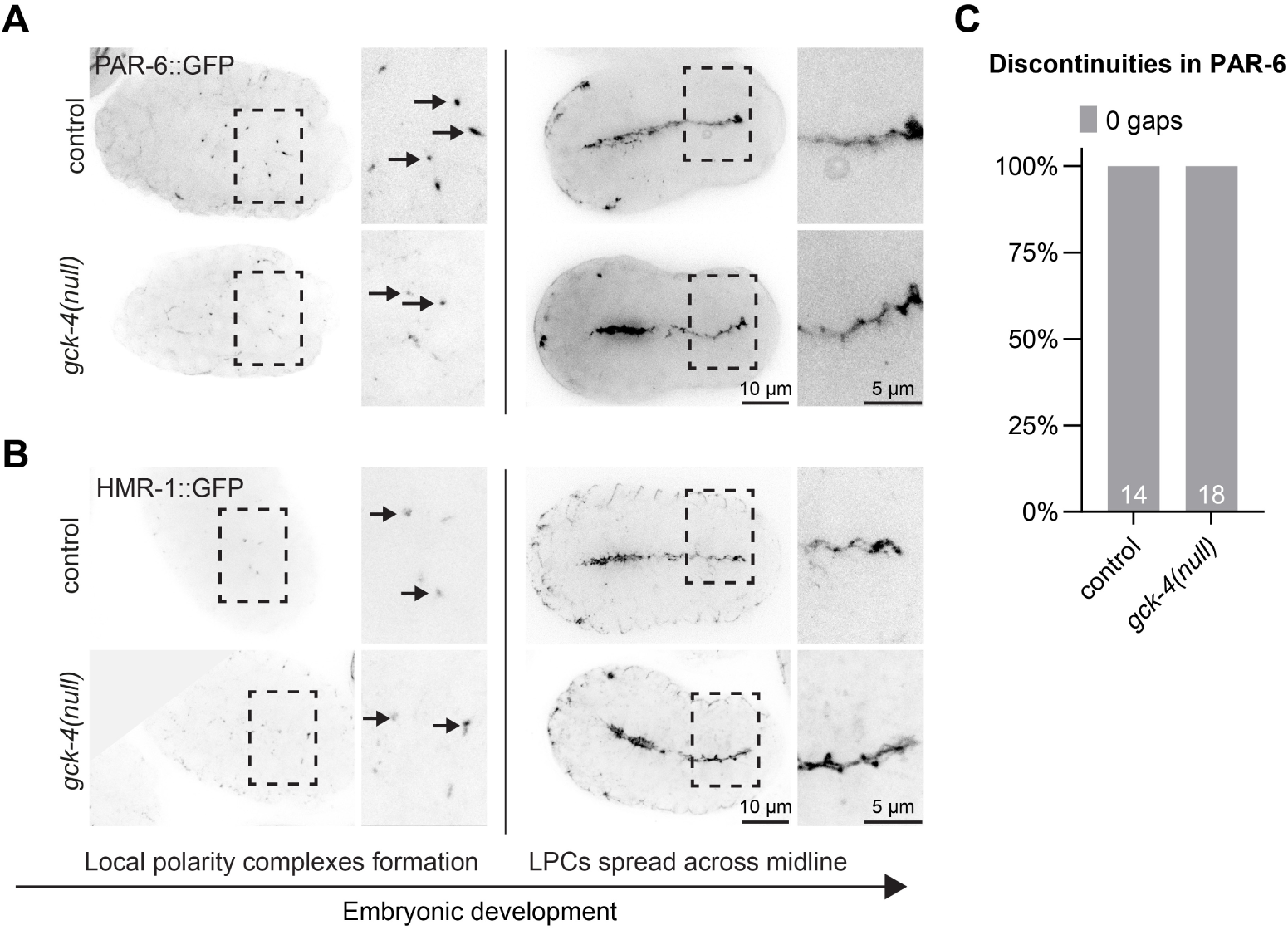
GCK-4 is not required for the establishment of intestinal epithelial polarity. (A, B) Maximum intensity projections of 3D stacks taken with a spinning disc confocal microscope showing the localization of PAR-6::GFP and HMR-1::GFP during polarity establishment in control and *gck-4(null)* embryos. Enlarged versions of the boxed regions are shown to the right of each panel. Arrows indicate individual local polarity complexes. Strains used: BOX251, BOX1129, BOX1034, and BOX1067. (C) Quantifications of gaps in PAR-6::GFP signal after localization along intestinal midline (right panels in A). Data are represented as a proportion of the total amount of embryos per genotype, with the total number per outcome indicated in the graph. Total number of embryos analyzed in order of samples in graph: 14 and 18. Data was analyzed with Fisher’s exact test; p value was not significant. Strains used: BOX251 and BOX1129.

### GCK-4 is essential for early intestinal lumen formation

As polarity establishment appeared unperturbed in *gck-4* mutant animals, we next examined the effects of GCK-4 loss on lumen formation. The intestinal lumen forms through cord-hollowing, initially forming small pockets that then connect to form a continuous lumen by the comma stage of development (Figure S3A). This is accompanied by segregation of apical, basolateral, and junctional components to their respective domains.

To examine the role of GCK-4 in these processes, we followed the distribution of the polarity proteins PAR-6 and LET-413 (Scribble), and the junctional proteins DLG-1 (Discs large) and HMR-1 using endogenous fluorescent protein fusions. By the 1.5-fold stage, embryos normally show a well-developed lumen decorated by PAR-6, surrounded by a ‘ladder-like’ junctional pattern of LET-413, DLG-1, and HMR-1 (Figure 4A, C). In contrast, *gck-4(null)* embryos showed a markedly reduced lumen diameter. Using the width of PAR-6::GFP as a proxy for lumen diameter, mutants displayed an average lumen width of 0.7 µm compared to 1.3 µm in controls (Figure 4B). The reduction in lumen diameter was also clearly visible in the localization pattern of DLG-1 and HMR-1, which additionally showed areas with no visible separation into two opposing periluminal junctions (the rails of the ladder) (Figure 4C). In areas with reduced lumen with but visible junction separation, DLG-1::mCherry fluorescence levels appeared normal, but HMR-1::GFP levels were elevated compared to controls (Figure 4D, S4A, B). In wider regions of the lumen, LET-413 was still excluded from the apical domain, consistent with our observations that overall polarity establishment is not affected by GCK-4 loss (Figure 4A; arrowheads).

**Figure 4:**
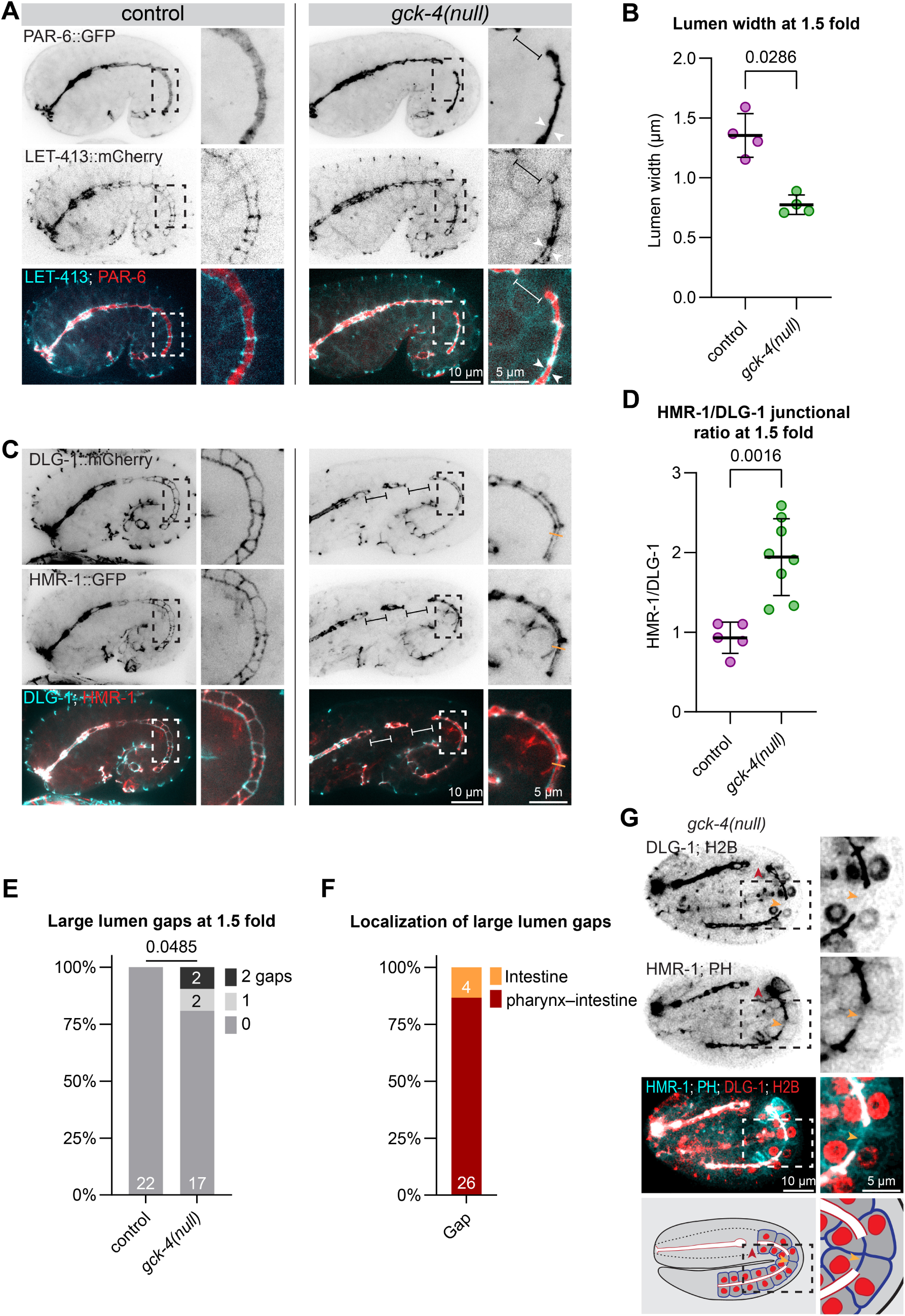
GCK-4 is essential for early intestinal lumen formation. (A) Maximum intensity projections of 3D stacks taken with a spinning disc confocal microscope showing the localization of PAR-6::GFP and LET-413::mCherry at the 1.5 fold developmental stage in control and *gck-4(null)* embryos. Enlarged versions of the boxed regions are shown to the right of each panel. White arrowheads show narrowed lumen and brackets show large midline gap in PAR-6 signal. Strains used: BOX251 and BOX1129. (B) Quantifications of lumen width in embryos as shown in (A). Lumen width was measured using the VasoMetrics tool in ImageJ and PAR-6::GFP signal. Data are represented as mean ± SD and analyzed with Mann-Whitney test; p value shown on graph. *n* = 4 for each condition. (C) Maximum intensity projections of 3D stacks taken with a spinning disc confocal microscope showing the localization of HMR-1::GFP and DLG-1::mCherry at the 1.5 fold developmental stage in control and *gck-4(null)* embryos. Brackets indicate large gaps in HMR-1::GFP and DLG-1::GFP signal. Enlarged versions of the boxed regions are shown to the right of each panel. Orange line exemplifies line used for quantifications in (D). Strains used: BOX1034 and BOX1067. (D) Quantifications of ratio between HMR-1::GFP and DLG-1::mCherry signal around lumen of embryos as shown in (C). Intensity values of HMR-1 and DLG-1 are shown in Figure S4A, B. Data are represented as mean ± SD and analyzed with Mann-Whitney test; p value shown on graph. Total number of embryos analyzed in order of samples in graph: 5 and 8. Strains used: BOX1034 and BOX1067. (E) Quantifications of large gaps in HMR-1::GFP and DLG-1::mCherry midline signal at 1.5 fold embryo stage. Examples of these gaps shown in (C). Data are represented as a proportion of the total amount of embryos per genotype, with the total number per outcome indicated in the graph. Data was analyzed with Fisher’s exact test; p value shown on graph. Total number of embryos analyzed in order of samples in graph: 22 and 21. Strains used: BOX1034 and BOX1067. (F) Quantification of the localization of gaps in HMR-1::GFP and DLG-1::mCherry midline signals as shown in (G). *N* = 30. Strain used: BOX1408. (G) Single focal plane spinning disc confocal microscopy image showing localization of DLG-1::mCherry, *Pges-1*::H2B::mCherry, *Pges-1*::GFP::PH and HMR-1::GFP in a 2-fold *gck-4(null)* embryo. Enlarged versions of the boxed regions are shown to the right of each panel. Red arrowhead indicates a large lumen gap between the pharynx and intestine, orange arrowhead indicates a gap within the intestine. A schematic of the phenotype is shown on the bottom. Strain used: BOX1408.

In addition to the narrowed lumen, we observed a large gap in the midline enrichment of PAR-6, HMR-1 and DLG-1 in ∼20% of embryos (Figure 4A, C, E). To better understand if such gaps represent a lack of apical domain or an intercalation defect resulting in a section of the intestine being formed by a single cell, we combined the junction markers with intestine specific expression of H2B::mCherry to mark nuclei and of the PH^PLC1δ^ domain fused to GFP to mark the membrane. The large lumen gaps were mostly located between the pharynx and first intestinal ring, indicating a role for GCK-4 in mediating the attachment of the posterior bulb of the pharynx to the anterior rings of the intestine (Figure 4F). In a small number of animals, we also observed larger gaps within the intestine itself. Within lumen gaps we still observed the GFP::PH membrane marker and gaps were flanked by two nuclei (Figure 4G). Moreover, the boundaries of the gaps did not align with the boundaries between intestinal rings, indicating that the gaps do not represent a specific intestinal ring failing to form a luminal domain.

Finally, we examined the luminal defects in more detail in larval stages. As in embryos, we observed both lumen constrictions and occasional larger gaps. The constrictions in larval stages fell into two categories. In one, junctions appeared ‘pinched’, but there was still a visible space between opposing junctions (orange arrowheads in Figure 5A). In the other, DLG-1 appeared as a single constricted point, completely separating two lumen pockets from each other (white arrows in Figure 5A). Constrictions were not limited to contact points between intestinal rings but appeared along the lengths of intestinal cells. The larger gaps, as in embryos, were predominantly present between intestine and pharynx, but also occasionally within the intestinal lumen (Figure 5A and Figure S5A). The fraction of L1 *gck-4(null)* larvae with large gaps was comparable to the number in 1.5-fold embryos, indicating that the luminal domain fragments during intestinal elongation (Figure 5B, Figure 4E). The occasional gaps within the intestinal lumen again did not match specific intestinal ring boundaries (Figure 5A), and we still observed LET-413 signal (Figure 5C), further confirming that the gaps within the lumen represent a failure to specify apical domain identity or maintain its continuity (Figure 5C).

**Figure 5:**
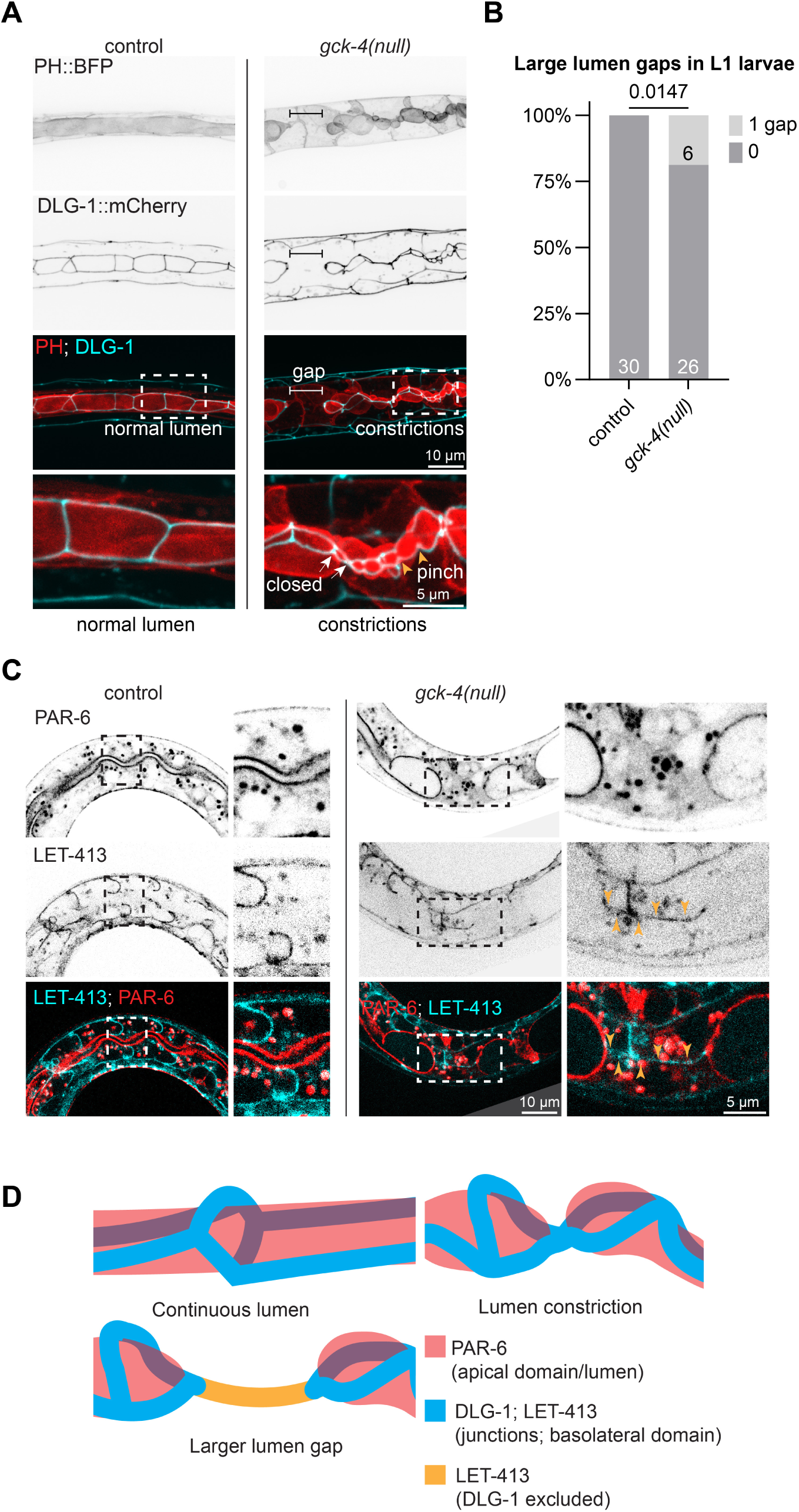
GCK-4 is essential for a continuous lumen in larvae. (A) Maximum intensity projections of 3D stacks taken with a spinning disc confocal microscope showing the localization of DLG-1::mCherry and *Pelt-2*::PH::mTagBFP in control and *gck-4(null)* L1 larvae. Bracket indicates large lumen gap. Enlarged versions of the boxed regions are shown under each panel. Strains used: BOX1034 and BOX1067. (B) Quantifications of large gaps in DLG-1::mCherry midline signal in L1 stage larvae as shown in (A). Total number of larvae analyzed in order of samples in graph: 30 and 32. Data are represented as a proportion of the total amount of larvae per genotype, with the total number per outcome indicated in the graph. Data was analyzed with Fisher’s exact test; p value shown on graph. Strains used: BOX1034 and BOX1067. (C) Maximum intensity projections of 3D stacks taken with a spinning disc confocal microscope showing the localization of PAR-6::GFP and LET-413::mCherry in control and *gck-4(null)* L1 larvae. Orange arrowheads indicate LET-413::mCherry localizing to a large lumen gap. Enlarged versions of the boxed regions are shown to the right of each panel. Strains used: BOX251 and BOX1129. (D) Schematic of continuous lumen found in control worms, and the two luminal defects observed in *gck-4(null)* larvae (lumen constriction and larger lumen gap).

In summary, the occluded intestine phenotype of *gck-4(null)* mutant larvae appears the result of a failure to properly expand the lumen and separate the junctions between opposing cells that form an intestinal ring (Figure 5D). Whether the lumen expansion defects are caused by a failure to separate junctions, or vice versa, cannot be distinguished. Additionally, in a subset of animals, the pharynx and intestine do not properly connect, or a larger section of the intestine fails to establish or maintain luminal domain identity.

### GCK-4 is required for apical enrichment of actin and ERM-1

The mammalian orthologs of GCK-4, LOK and SLK, have emerged as key regulators of the apical actin network (Viswanatha et al., 2012; Zaman et al., 2021). In epithelial cells, LOK and SLK activity is essential for microvillus formation, while their loss induces abnormal apical actomyosin bundles and perturbs cell–cell junction organization (Viswanatha et al., 2012; Zaman et al., 2021). Our ultrastructural analysis similarly shows that the loss of GCK-4 severely impacts microvillus formation. These observations prompted us to investigate whether GCK-4 organizes the apical actin network and thereby contributes to lumen formation.

To visualize apical actin organization, we used a marker strain expressing YFP fused to ACT-5, the predominant intestinal actin isoform in *C. elegans*. In control embryos, luminal YFP::ACT-5 was detected by the bean stage and became progressively enriched during embryonic development (Figure 6A, C). In larvae, YFP::ACT-5 remained enriched at the apical membrane and in the actin-rich microvillar brush border (Figure S6A). In *gck-4(null)* embryos, enrichment of ACT-5 at the apical domain was delayed until the 2-fold stage, and overall levels were sharply reduced (Figure 6A, C). Apical actin levels remained strongly reduced in L1-stage *gck-4* animals, both using YFP::ACT-5 and LifeAct::mRuby as an alternative marker (Figure S6A–D).

**Figure 6:**
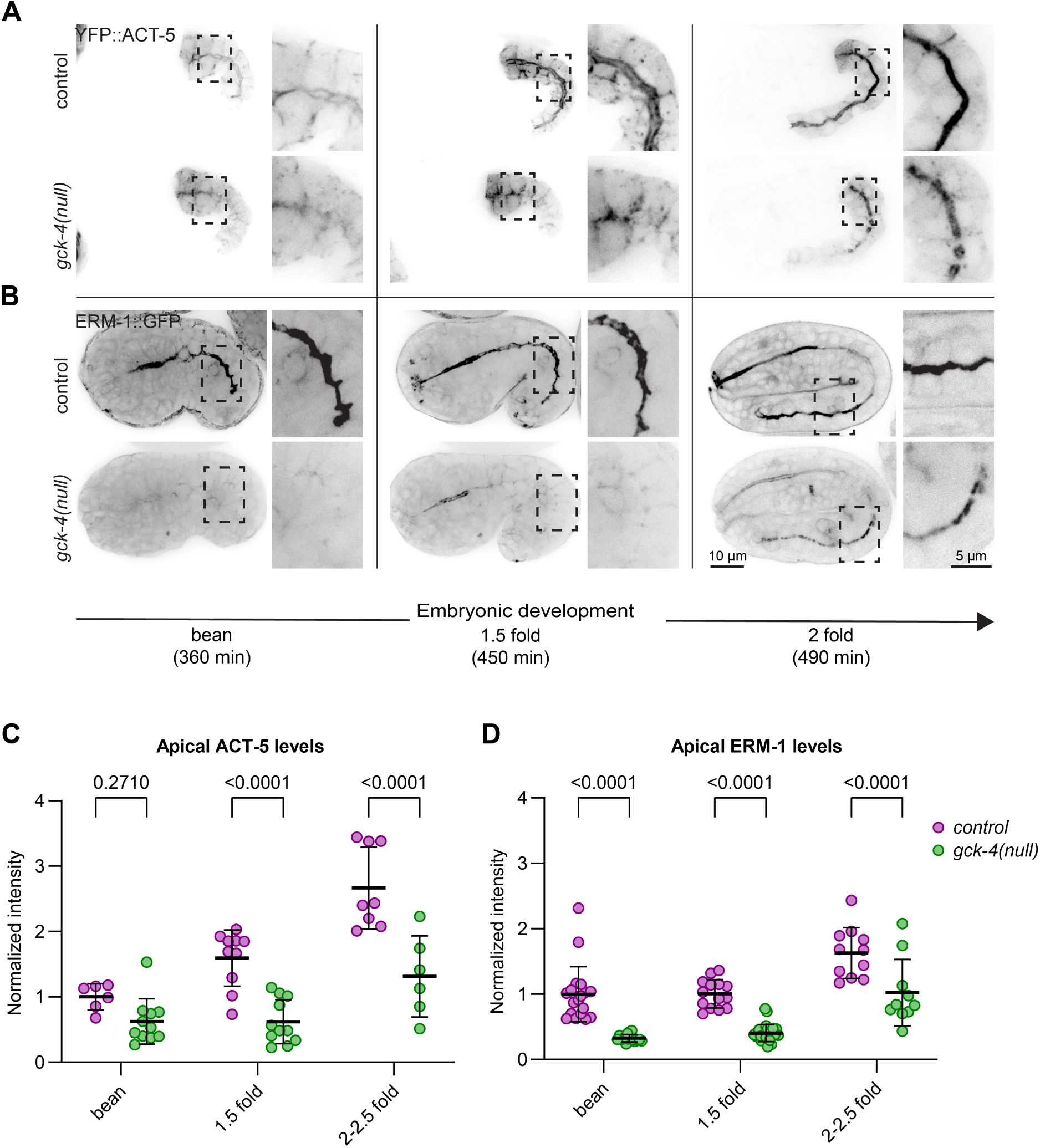
GCK-4 is required for apical enrichment of actin and ERM-1. (A, B) Maximum intensity projections of 3D stacks (bean and 1.5-fold) and single focal plane (2-fold) taken with a spinning disc confocal microscope showing the localization of *pGES-1*::YFP::ACT-5 and ERM-1::GFP during intestinal development. Enlarged versions of the boxed regions are shown to the right of each panel. Strains used: BOX1031, BOX1059, BOX1030, and BOX1065. (C) Quantifications of apical YFP::ACT-5 levels during intestinal development. Data are represented as mean ± SD and analyzed with Mann-Whitney test; p value shown on graph. Total number of embryos analyzed in order of samples in graph: 6, 11, 10, 11, 8 and 6. Strains used: BOX1031 and BOX1059. (D) Quantifications of apical ERM-1::GFP levels during intestinal development. Data are represented as mean line ± SD and analyzed with Mann-Whitney test; p value shown on graph. Total number of embryos analyzed in order of samples in graph: 19, 15, 14, 25, 11 and 10. Strains used: BOX1030 and BOX1065.

LOK and SLK are thought to regulate the apical actomyosin network in large part through their regulation of Ezrin-Radixin-Moesin (ERM) proteins (Pelaseyed et al., 2017; Viswanatha et al., 2012; Zaman et al., 2021). ERM proteins serve as molecular linkers between the plasma membrane and the actin cytoskeleton and can organize specialized membrane domains (Fehon et al., 2010). The sole *C. elegans* ortholog, ERM-1, is a key actin regulator in the intestine, and its loss results in morphological and luminal defects similar to those observed in *gck-4* mutants (Göbel et al., 2004; Ramalho et al., 2020; Sepers et al., 2022). We therefore next investigated the localization of ERM-1.

In control animals, ERM-1::GFP accumulated at the nascent lumen by the bean stage, and remained enriched at the luminal domain throughout embryogenesis and larval stages (Figure 6B, S6E, F). To more precisely determine the timing of ERM-1 recruitment, we analyzed the localization of ERM-1::mCherry relative to that of PKC-3::GFP, a component of the apical PAR complex and of local polarity complexes (Figure S6G). ERM-1 was recruited to the midline after the arrival of PKC-3, indicating that ERM-1 is not a component of the local polarity complexes, but is recruited to the midline immediately before or during lumen initiation. In contrast, in *gck-4(null)* mutant embryos ERM-1::GFP failed to enrich at the midline during the bean and 1.5 fold stages, when the lumen normally fully forms (Figure 6B, 6D). Only at the 2-fold embryonic stage did ERM-1::GFP begin to accumulate apically, but at reduced intensity compared to controls. In *gck-4(null)* larvae, ERM-1::GFP remained enriched at the luminal domain, but at reduced levels compared to controls (Figure S6E, F).

ACT-5 and ERM-1 thus show similar defects in *gck-4* mutant embryos: enrichment at the luminal domain occurs much later and at reduced levels than in controls, and these defects persist to the L1 stage, at which point mutants arrest development. The reduced ACT-5 and ERM-1 levels could contribute to the impaired microvilli formation or, alternatively, be an indirect consequence of it. The timing of the defects argues for the former: ACT-5 and ERM-1 recruitment defects are already visible at the bean stage, while little microvilli formation is observed before the 2-fold stage (Bidaud-Meynard et al., 2021). Together, these results indicate that GCK-4 is required for the proper apical recruitment of ACT-5 and ERM-1 and for organization of the apical actin cytoskeleton, following polarity establishment and during lumen formation. Their reduced levels likely contribute to the lumen and microvilli defects.

### GCK-4 is not the major ERM-1 T544 kinase in the intestine

The mammalian kinases LOK and SLK regulate microvilli formation via activation of ERM proteins through phosphorylation of a highly conserved regulatory C-terminal threonine (Pelaseyed et al., 2017; Viswanatha et al., 2012; Zaman et al., 2021). Moreover, LOK/SLK and their *Drosophila* ortholog Slik are proposed to be the main or only kinases that phosphorylate ERM proteins (Hipfner et al., 2004; Hughes and Fehon, 2006; Pelaseyed et al., 2017; Viswanatha et al., 2012; Zaman et al., 2021). In comparison with *gck-4* mutants, animals expressing non-phosphorylatable ERM-1[T544A] have less severe intestinal defects that resolve by the L1 stage, and are more viable (Ramalho et al., 2020). This makes it unlikely that the *gck-4* phenotypes can be fully explained by a lack of ERM-1 phosphorylation (Ramalho et al., 2020). Nevertheless, we sought to verify whether the regulatory relationship between LOK/SLK and ERM proteins is conserved in *C. elegans*. To test whether GCK-4 phosphorylates ERM-1 at T544, we made use of an antibody specific to ERM proteins phosphorylated at this residue (α-pERM). The antibody was raised against an Ezrin polypeptide which is also fully conserved in *C. elegans* ERM-1, and we previously demonstrated its specificity by showing that it does not cross-react with a T544A mutant variant of ERM-1, and that immunostaining of fixed embryos is abolished by treatment with a phosphatase (Sepers et al., 2022).

To test whether loss of GCK-4 affects the phosphorylation status of ERM-1, we analyzed protein lysates prepared from newly-hatched larvae by western blot. As the intestine is the major ERM-1 expressing tissue at this stage, the majority of western blot signal will represent intestinal ERM-1. For this analysis, we made use of homozygously maintained *gck-4(null)* and *gck-4(K64R)::mCherry::AID* kinase dead mutants, to eliminate the possibility of phosphorylation by perduring maternal wild-type *gck-4* product. To control for loading, we performed the analyses in strains endogenously expressing ERM-1::GFP, which is readily detected with an anti-GFP antibody. As expected, ERM-1::GFP was phosphorylated in control worms, whereas no phosphorylation was detected in *erm-1(T544A)* mutants (Figure 7A). As an extra control for antibody specificity we also performed lambda phosphatase treatment on membranes after protein transfer. ERM-1 T544 phosphorylation was no longer detectable, whereas ERM-1::GFP detection levels remained unchanged, confirming that the antibody is specific for phosphorylated T544 on western blot (Figure S7A). Unexpectedly, phosphorylation levels in *gck-4(null)* and *gck-4(K64R)::mCherry::AID\** kinase dead mutants only showed a reduction of ∼40% compared to control, rather than a complete loss.

**Figure 7:**
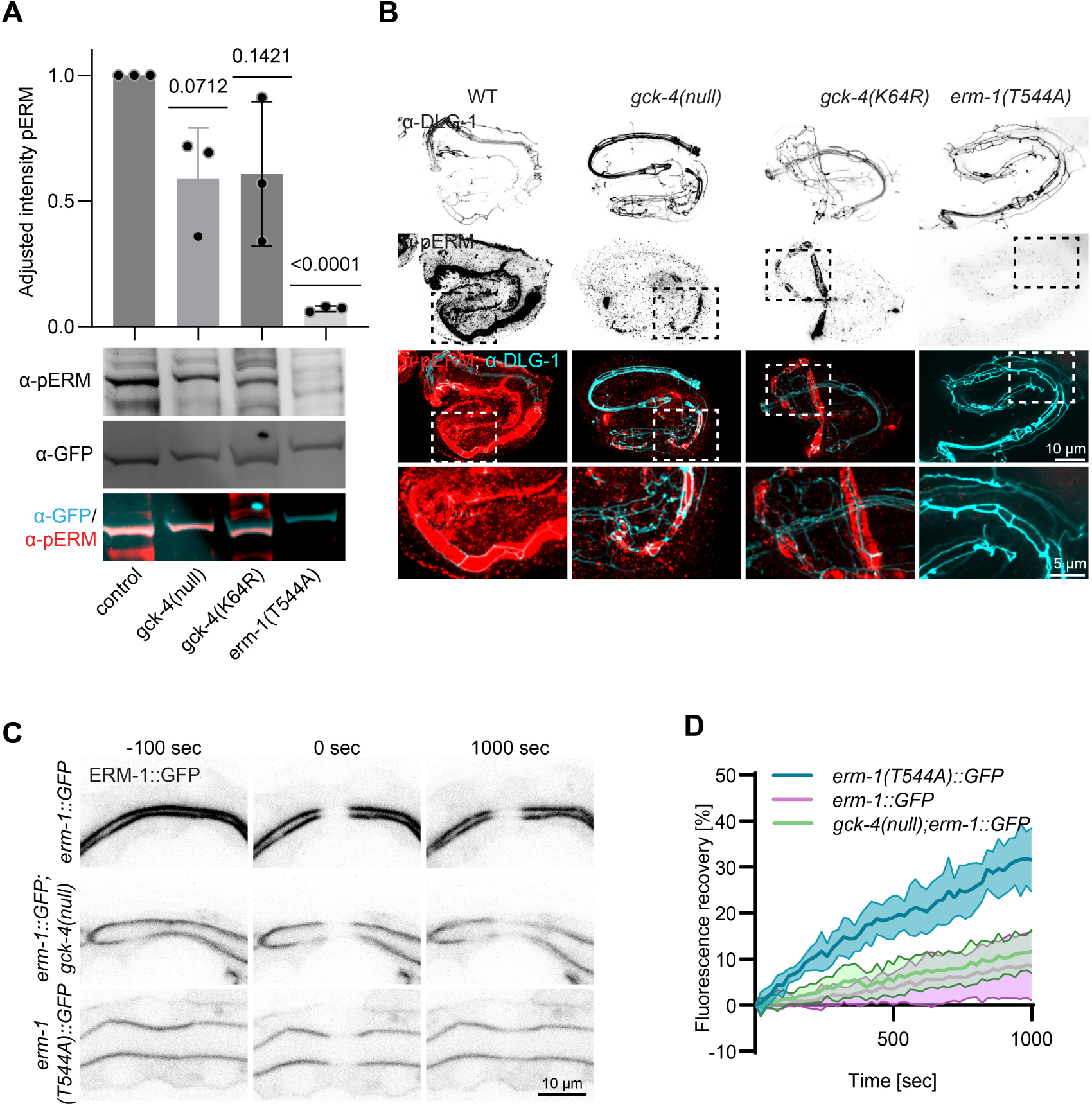
GCK-4 regulation of the actin cytoskeleton is independent of ERM-1 T544 phosphorylation. (A) Levels of ERM-1 T544 phosphorylation in freshly hatched L1 larvae assessed by western blot. Data are represented as mean ± SD and analyzed with a one sample t-test with a theoretical mean of 1 for each condition; p value shown on graph. α-pERM = phosphorylated ERM-1 T544, α-GFP = total ERM-1::GFP, *N* = 3. Data points in graph represent individual quantifications. Example blot shown below graph and all blots shown in Supplementary Figure S7C. Strains used: BOX1030, BOX1065, BOX1409 and BOX369. (B) Representative images of fixed embryos stained with antibodies recognizing the junctional protein DLG-1 (α-DLG) and phosphorylated ERM-1 (α-pERM), imaged with spinning disc confocal microscopy. Enlarged versions of the boxed regions are shown under each panel. Strains used: CGC1, BOX1036, BOX1402 and BOX165. (C) Representative single focal plane views taken with a spinning disc confocal microscope showing the localization and intensity of ERM-1::GFP before and after FRAP bleaching in the intestine of L1 larvae of indicated genotypes. (D) FRAP analysis of apical ERM-1::GFP in the intestine of L1 larvae. Graph shows the fluorescence intensity of ERM-1 in the photobleached region at the apical intestinal domain during recovery. Data are represented as mean ± SD. Number of animals per genotype: *erm-1::GFP* = 6, *gck-4(null);erm-1::GFP* = 8, *erm-1(T544A)::GFP* = 7. Strains used: BOX1030, BOX1065 and BOX369.

The western blot results cannot rule out that GCK-4 loss specifically affects ERM-1 phosphorylation in the intestine or during earlier embryonic stages. Therefore, we performed immunostaining for phosphorylated ERM-1 in control, *gck-4(null), gck-4(K64R)* and *erm-1(T544A)* embryos (Figure 7B). In all conditions except *erm-1(T544A)*, we detected phosphorylated ERM-1 at the intestinal midline from the 2-fold stage on. Thus, from the moment in development that ERM-1 enrichment becomes apparent in *gck-4* mutant embryos, we also detect phosphorylation of ERM-1 on T544. While a short time window remains during which ERM-1 is not enriched at the midline, and its phosphorylation status cannot be assessed, these results strongly argue that GCK-4 is not the main regulator of ERM-1 T544 phosphorylation in the intestine of *C. elegans*.

We next investigated ERM-1::GFP dynamics using fluorescence recovery after photobleaching (FRAP) in L1 larvae. As we previously found, ERM-1[T544A] exhibited much faster recovery than control ERM-1::GFP (Figure 7C, D). In contrast, ERM-1::GFP recovery in *gck-4(null)* mutants was indistinguishable from that in *gck-4* wild-type animals, consistent with our observation that ERM-1::GFP is still phosphorylated in *gck-4(null)* larvae.

Finally, we investigated whether expression of phosphorylation mimicking ERM-1[T544D] could rescue the phenotype of *gck-4* mutants (Figure S7B). While we did observe a reduction in larval lethality, we obtained a similar level of rescuing activity by expressing the non-phosphorylatable ERM-1[T544A] variant. This result indicates that the partial rescue of the *gck-4* phenotype is due to a previously unrecognized increase in ERM-1 activity resulting from mutating the T544 site, rather than due to mimicking of the T544 phosphorylation state.

Taken together, our data indicate that, while GCK-4 regulates apical accumulation of ERM-1 during lumen formation, this regulation is not mediated, or is only partially mediated, via phosphorylation of its regulatory C-terminal threonine. As our analysis of the kinase dead *gck-4(K64R)* mutant indicates that kinase activity is essential for GCK-4 function, this implies that GCK-4 has other targets.

## Discussion

In this study, we identified GCK-4, the *C. elegans* ortholog of the Ste20p-family kinases LOK and SLK, as an essential regulator of lumen formation in the intestine. Loss of GCK-4 resulted in a constricted and cystic intestine with severe defects in microvilli formation, resulting in ∼98% lethality. While a seemingly continuous apical domain was established during intestinal polarization, larger gaps in the midline started to form during intestinal elongation in a subset of embryos, particularly between the pharynx and first intestinal ring. Thus, GCK-4 is essential for intestinal lumen formation and protects lumen continuity during elongation.

The lumen defects we observed were accompanied by severe defects in the establishment of the normally highly prominent apical actin cytoskeleton. During early steps in lumen formation, we observed little apical enrichment of actin in *gck-4* mutants. From the 2-fold stage on, actin was enriched at the luminal domain, but levels remained lower than in control animals. The apical cytoskeleton plays essential roles in organizing the luminal domain, and apical enrichment of F-actin is a feature of most, if not all multicellular epithelial lumens, including MDCK cysts, mammalian enterocytes, *Drosophila* embryonic tissues and endothelial cells (Hirokawa et al., 1982; Kondrychyn et al., 2020; Martin-Belmonte et al., 2007; Massarwa et al., 2009). The loss of apical actin accumulation is likely to contribute greatly to the *gck-4* phenotype. Nevertheless, changes in actin levels alone may not explain the full *gck-4* phenotype, as we previously observed apical actin accumulation defects in mutants expressing non-phosphorylatable ERM-1[T544A], which have far less severe intestinal defects (Ramalho et al., 2020). Thus, apical enrichment defects in *gck-4* mutants may be accompanied by additional functional defects in the apical cytoskeleton or other components of the luminal domain.

Apical junctions were also strongly affected by loss of GCK-4. During embryonic development, opposing junctions along the lumen failed to fully separate, and in areas showed no detectable separation at all. In these regions HMR-1 (E-cadherin) levels, but not DLG-1 levels, were enriched. In larvae, junctions continued to show defects in the form of constriction. Ultrastructurally, junctions were elongated and the electron-dense material in EM images was irregularly clustered rather than evenly distributed. The failure to separate junctions and the failure to expand the lumen may be caused by the same underlying defect. The apical actin cytoskeleton and junctions are mechanically connected: junctions anchor the cortical actin network and transmit actin-generated tension between cells, and actin organization and contractility in turn shape the organization of adherens junctions (Cavey and Lecuit, 2009; Mège et al., 2006). The reduced apical actin we observe in *gck-4* mutants could therefore contribute to the junctional defects, though GCK-4 may also act on the junctions and the cytoskeleton through separate targets. Interestingly, defects in the organization of apical junctions and junctional actin were also found in LOK/SLK and ezrin/radixin knockout cells (Zaman et al., 2021), and we previously observed increased HRM-1 levels in the rare constrictions present in ERM-1[T544A] L1 larvae (Ramalho et al., 2020). Hence, the junctional defects may also be caused at least in part by the reduced levels of apical ERM-1 present in *gck-4* mutant animals.

Based on the roles of GCK-4 orthologs in other systems, we hypothesized that GCK-4 regulates lumen formation via phosphorylation of ERM-1, the single *C. elegans* member of the Ezrin/Radixin/Moesin (ERM) family. ERM proteins are critical regulators of microvilli formation due to their ability to link the microvillar actin core to the plasma membrane (Fehon et al., 2010). Indeed, loss of ERM-1 in *C. elegans* leads to cystic lumens with reduced microvilli formation (Göbel et al., 2004), phenotypes that resemble those caused by GCK-4 loss. Moreover, in *Drosophila* and mammalian cells, GCK-4 orthologs have been found to regulate ERM proteins via phosphorylation of their conserved C-terminal threonine residues. In *Drosophila* the kinase Slik regulates epithelial integrity in imaginal disks through regulation of Moesin phosphorylation (Hipfner et al., 2004; Hughes and Fehon, 2006), and similarly controls Moesin activation at the mitotic cell cortex in cultured S2 cells (Carreno et al., 2008; Kunda et al., 2008). In mammalian epithelial cells, particularly Jeg-3 cells, LOK and SLK are thought to be the predominant or only kinases that phosphorylate ERM proteins, and ERM proteins appear to be the only target involved in the phenotypes of LOK^−/−^ SLK^−/−^ cells (Lombardo et al., 2024; Pelaseyed et al., 2017; Viswanatha et al., 2012; Zaman et al., 2021). However, our results are not consistent with GCK-4 acting solely through phosphorylation of ERM-1, as *gck-4* null and kinase-dead mutants retain substantial ERM-1 phosphorylation, detected both by western blot in larvae and by immunostaining in embryos. Moreover, we previously demonstrated that a non-phosphorylatable ERM-1[T544A] mutant has only mild defects in lumen establishment or microvilli formation (Ramalho et al., 2020). Thus, loss of ERM-1 phosphorylation alone cannot explain the severe phenotypes observed in *gck-4* mutant animals.

We considered the possibility that the phosphorylated ERM antibody recognizes unphosphorylated ERM-1, as pERM antibodies have been reported to have some reactivity to unphosphorylated ezrin (Pelaseyed et al., 2017). We previously demonstrated that the pERM antibody does not stain embryos expressing ERM-1(T544A) nor embryos expressing ERM-1 with an unaltered T544 residue but treated with a phosphatase (Sepers et al., 2022). We now show that the antibody also does not recognize ERM-1(T544A) on western blot nor unaltered ERM-1(T544) on a phosphatase-treated western blot. Hence, the pERM antibody appears highly specific under the conditions used here.

We also observed partial rescue of larval lethality in *gck-4* mutants by expression of the phosphorylation mimicking ERM-1[T544D] variant. However, expression of non-phosphorylatable ERM-1[T544A] rescued viability to the same extent. These results indicate that mutation of the T544 residue in itself, rather than the change to an aspartic acid, is responsible for the rescuing activity, for example by increasing the dynamics of ERM-1.

While GCK-4 does not appear to be the major ERM-1 kinase in the *C. elegans* intestine, it remains possible that ERM-1 phosphorylation defects contribute to the *gck-4* phenotype. First, while loss of GCK-4 does not abolish ERM-1 phosphorylation, pERM-1 levels are reduced by ∼40% on western blots of *gck-4* mutant larvae. This reduction could indicate a contribution of GCK-4 to ERM-1 phosphorylation, but it could also be an indirect effect. For example, loss of GCK-4 reduces apical levels of ERM-1, which in turn could reduce phosphorylation by an apical kinase. Alternatively, loss of *gck-4* might affect the localization or activity of ERM-1 kinase(s). Second, there is a short window of time in early intestinal development where we cannot determine the ERM-1 phosphorylation status in *gck-4* mutants due to the delay in apical enrichment of ERM-1. It remains possible therefore that GCK-4 plays an important role in ERM-1 phosphorylation during the early stages of intestinal development.

Despite the caveats above, our data support that GCK-4 likely has other targets than ERM-1 that are important in lumen formation. One interpretation of these results is that, while the essentiality of ERM-1 and GCK-4 proteins is conserved, regulatory aspects of these proteins have evolved differently in *C. elegans* than in *Drosophila* and mammals. An alternative explanation is that the importance of ERM protein phosphorylation and the key GCK-4/LOK/SLK targets vary between organisms or tissue types. For example, the LOK/SLK – ERM pathway is extensively studied in Jeg-3 cells, which are human choriocarcinoma cells derived from placenta (Kohler et al., 1971). However, in CACO-2 cells, a human colorectal adenocarcinoma cell line, PKCι has been reported to be the kinase responsible for activating ezrin in the context of brush border formation (Wald et al., 2008). Whether LOK and SLK are important ezrin kinases in this cell type is difficult to assess as Caco-2 cells lacking LOK and SLK cannot be maintained (Zaman et al., 2021). Similarly, while the importance of ERM phosphorylation is widely reported, there are indications that its importance is not universal. For example, in *Drosophila,* a non-phosphorylatable form of Moesin was able to significantly rescue the viability of *Moe* mutant flies and can mediate normal wing-disc epithelium formation (Roch et al., 2010). In mammalian cell lines, phosphorylation of ERM proteins was found to be dispensable for the formation of microvilli-like structures in A-431 epidermal epithelial cells and MDCK II canine kidney cells (Yonemura et al., 2002). In *C. elegans*, development of the mature brush border progresses in stages, as components of the brush border are highly dynamic during embryonic development, but become very stable in larval stages (Bidaud-Meynard et al., 2021). One possibility is that the *C. elegans* intestine, developing in the context of a full organism, represents a more complex system where multiple regulatory mechanisms are at play.

Regardless of the reason for the observed differences, two important questions will need to be addressed in future experiments. The first is the nature of the ERM-1 kinase. A number of candidate kinases have been proposed to phosphorylate ERM proteins in addition to LOK/SLK, including NCK interacting kinase (NIK) (Baumgartner et al., 2006), Leucine-rich repeat protein kinase 2 (LRRK2) (Parisiadou et al., 2009), CDK5 (Yang and Hinds, 2003), Protein Kinase Cθ (Pietromonaco et al., 1998) and Protein Kinase Cα (Ng et al., 2001), Macrophage stimulating 4 (Mst4) (Gloerich et al., 2012; ten Klooster et al., 2009), JNK (Pan et al., 2013), and Rho-associated kinase (ROCK) (Haas et al., 2007; Matsui et al., 1998; Tran Quang et al., 2000). Our RNAi screen did not identify strong intestinal defects for the kinases GCK-1 (Mst4), MIG-15 (NIK), LET-502 (ROCK), PKC-2 (Protein Kinase Cα), TPA-1 (Protein Kinase Cθ), or CDK-5. However, RNAi depletion may be incomplete and for *let-502* we had to rely on partial depletion to bypass earlier embryonic lethality. Revisiting these and other candidates with intestine-specific depletion or knockout approaches may therefore still reveal roles in ERM-1 phosphorylation.

A second important question to address is the identity of the GCK-4 target(s). The phenotypes of our *gck-4* kinase-dead allele are nearly as severe as those caused by complete deletion of the *gck-4* locus, indicating that kinase activity accounts for most of GCK-4 function. The slightly increased viability of the kinase-dead mutant suggest at most a minor kinase-independent contribution, consistent with the partly catalysis-independent functions previously reported for GCK-4 orthologs (Pelaseyed et al., 2017; Rambaud et al., 2025). Identifying the relevant phosphorylation targets therefore remains important. Additional targets have been identified for Slik and SLK. These include Talin, which is phosphorylated by *Drosophila* Slik to mediate maintenance of muscle attachment (Katzemich et al., 2019); Paxillin, whose phosphorylation by SLK mediates focal adhesion turnover in mammalian cells (Quizi et al., 2013); and a minor isoform of the p150^Glued^ subunit of dynactin, which was reported to localize to the centrosome upon phosphorylation by SLK (Zhapparova et al., 2013). The expression of the *C. elegans* Talin and Paxillin orthologs TLN-1 and PXL-1 is limited to skeletal muscle (both proteins), somatic gonad (TLN-1), and pharyngeal muscle (PXL-1) (García-Alvarez et al., 2003; Sadeghian et al., 2023; Warner et al., 2011). Moreover, their loss-of-function phenotypes are not consistent with intestinal defects related to GCK-4 loss. Gene expression analyses indicate that the p150^Glued^ ortholog DNC-1 is expressed in the intestine (Ghaddar et al., 2023; Sternberg et al., 2024), but its role in this tissue has not been investigated and loss of *dnc-1* is early embryonic lethal (Skop and White, 1998). Identification of the GCK-4 targets will therefore likely require unbiased approaches such as phosphoproteomics.

In summary, our experiments identify GCK-4 as an essential regulator of lumen formation in the *C. elegans* intestine, and highlight potential complexities in the regulation of the intestinal brush border that may be interesting to investigate in mammalian systems.

## Material and Methods

### *C. elegans* strains and culture conditions

*C. elegans* strains were cultured under standard conditions (Brenner, 1974). Only hermaphrodites were used, and all experiments were performed with animals grown at 15 °C or 20 °C on standard Nematode Growth Medium (NGM) agar plates seeded with OP50 *Escherichia coli*, unless indicated otherwise. Table 2 contains a list of all the strains used.

**Table 2.**
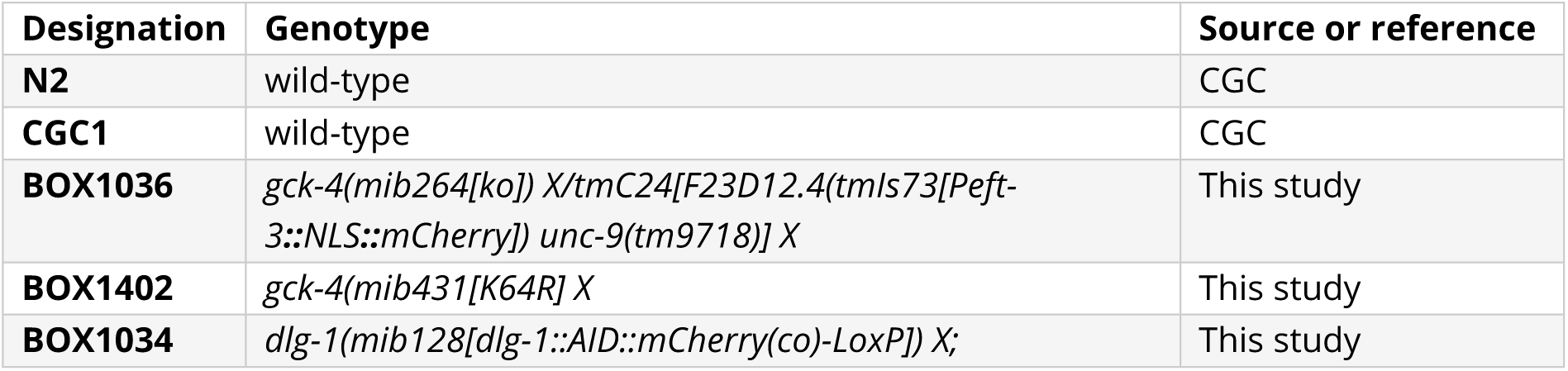
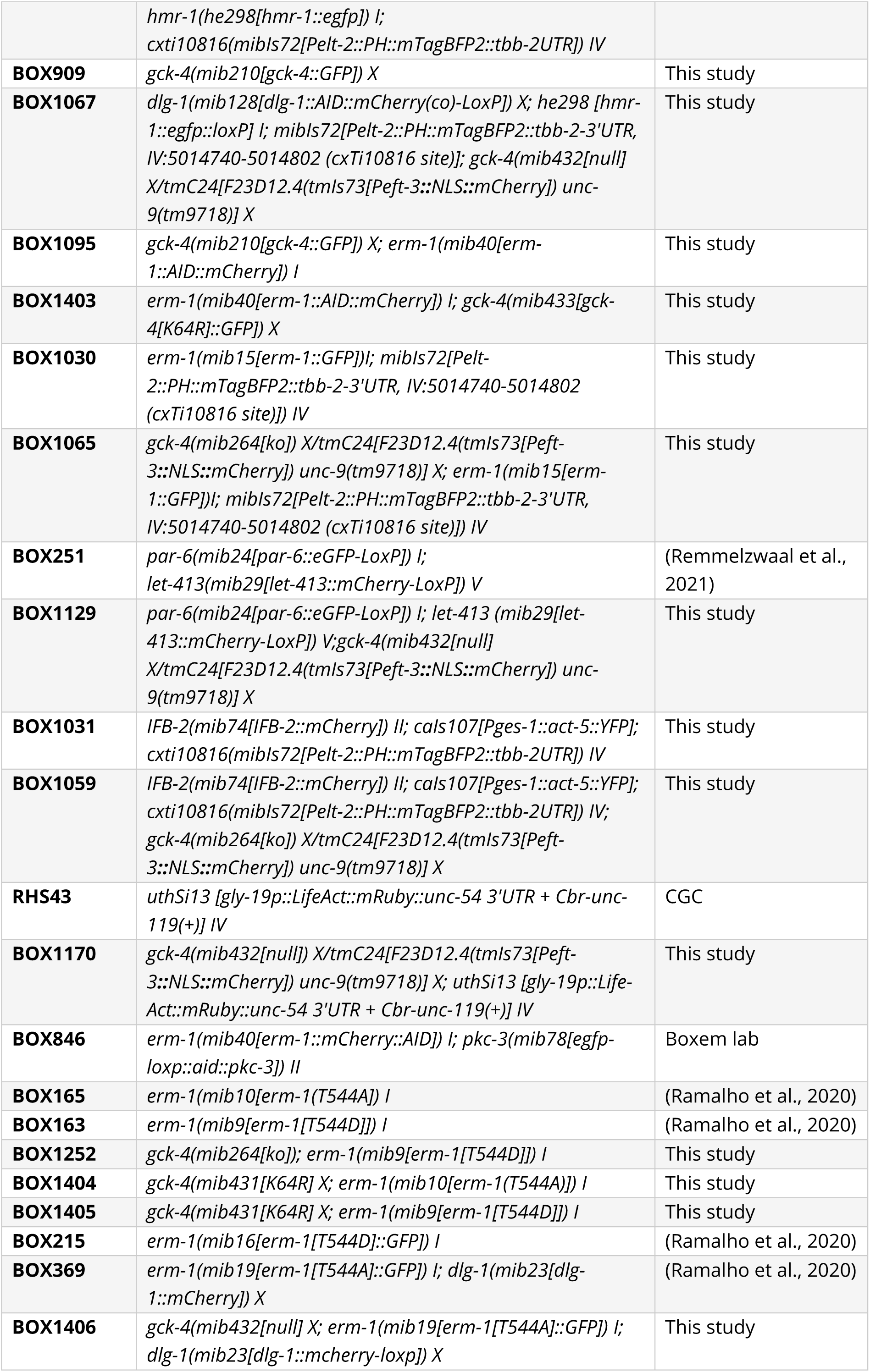
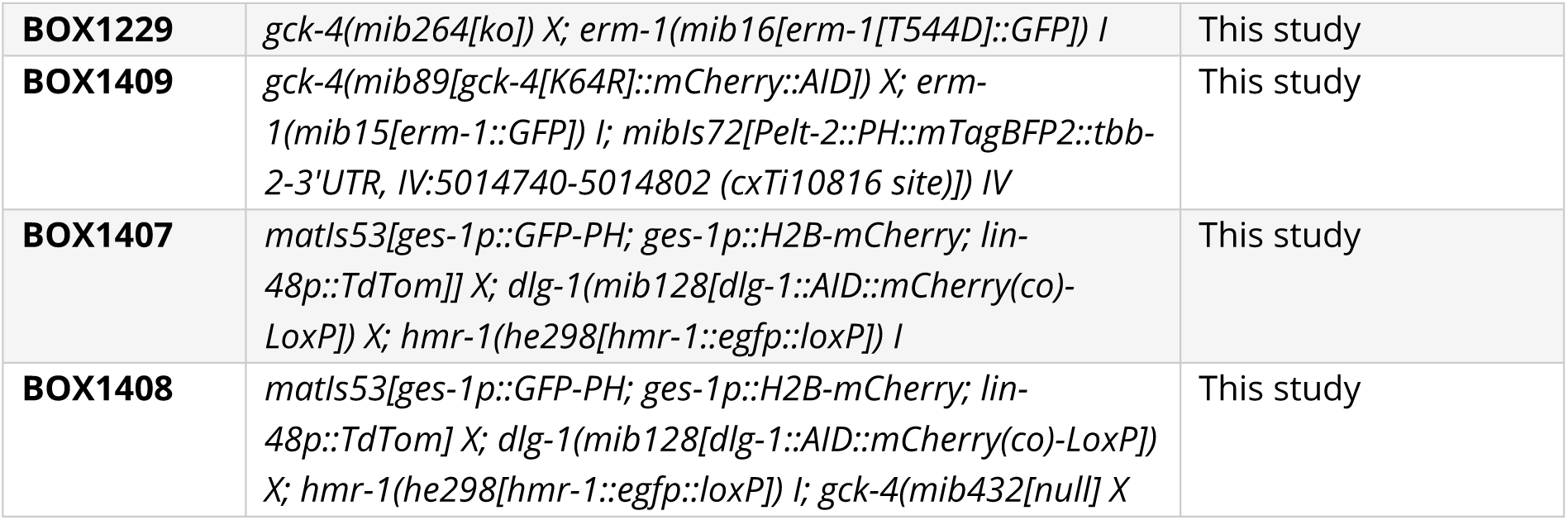
List of *C. elegans* strains used in the study.

### CRISPR/Cas9 genome editing

All alleles were made using homology-directed repair of CRISPR/Cas9-induced DNA double-strand breaks, for which the reagents were micro-injected into the gonad of young adults. If appropriate, repair templates included silent mutations to prevent recutting of repaired loci by Cas9. To verify the edits, the insertion sites were PCR amplified and sequenced by Sanger sequencing. The *gck*-*4::mCherry::AID\** allele was made using plasmid based expression of Cas9, sgRNA, and a plasmid repair template containing a self-excising cassette for selection of candidate integrants (Dickinson et al., 2015). The *gck*-*4[K64R]::mCherry::AID\** allele was generated by introducing the K64R mutation in the existing *gck*-*4::mCherry::AID\** allele, using Cas9 and sgRNA plasmids and a DNA oligo repair template with ∼35 bp homology arms. All other edits were made using Alt-R CRISPR/Cas9 reagents (IDT), as previously described (Ghanta and Mello, 2020). The repair template for the *gck*-*4::GFP* allele was amplified using primers with 5’ SP9 modifications (IDT) from the PJJS001 (Addgene #188324) plasmid. DNA oligos were used as repair templates for the *gck-4(null), gck-4(K64R)* and *erm-1* phosphomutant alleles. Table 3 contains a list of all CRISPR/Cas9 edits made with the respective oligo sequences (IDT). Supplementary file 1 contains Genbank format sequence files of edited loci (pre- and post-editing) annotated with reagent information.

**Table 3.**
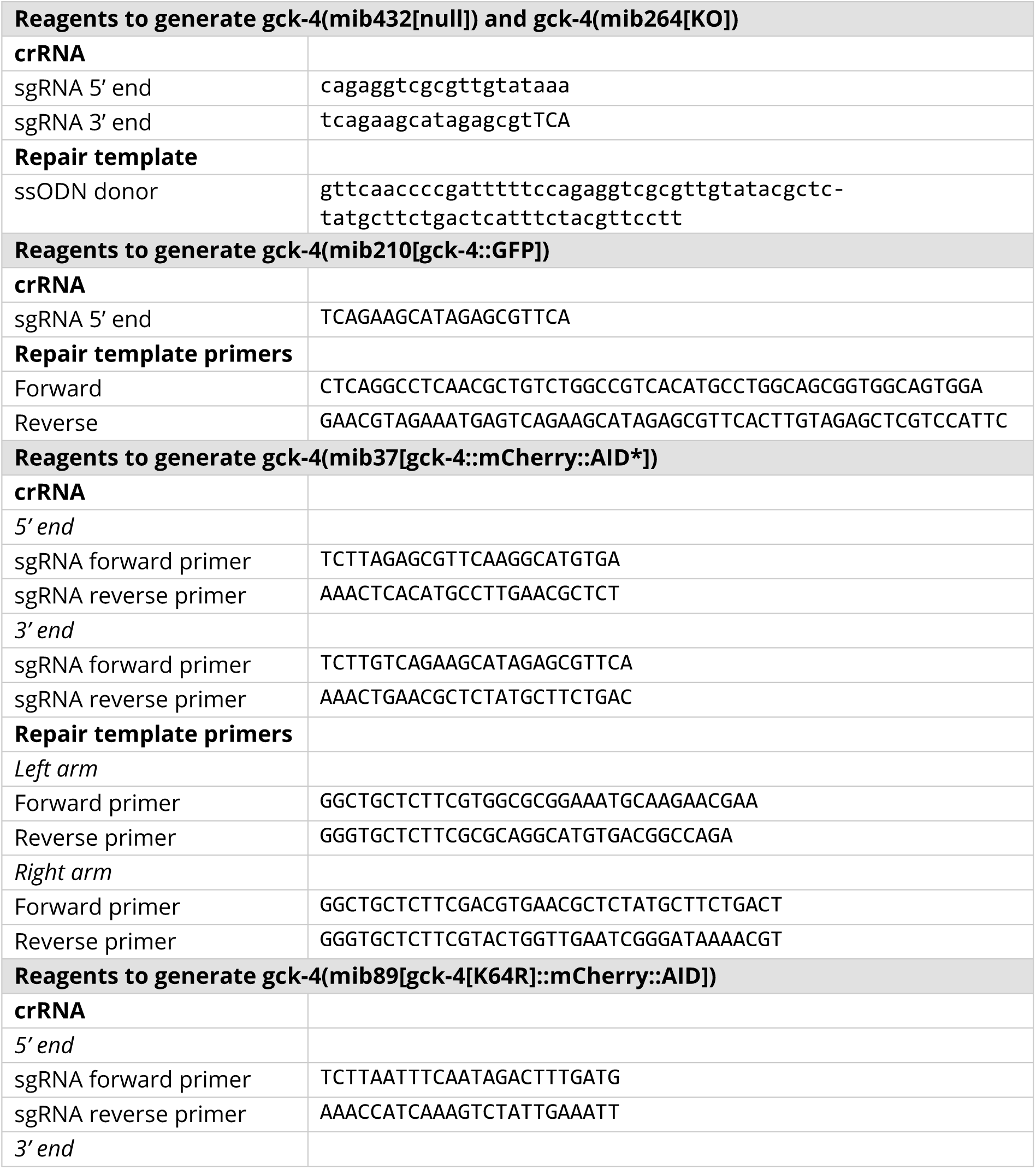
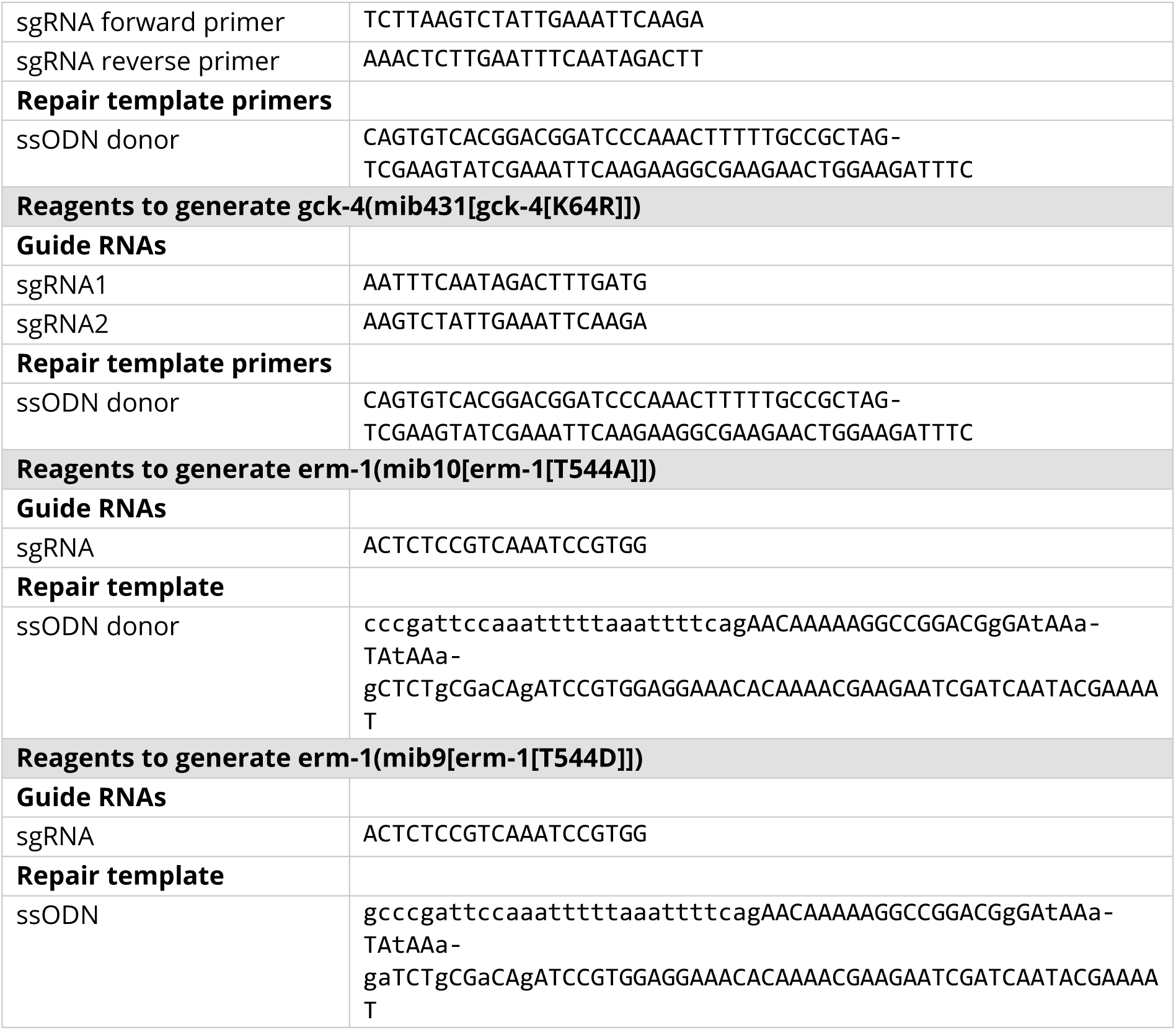
List of guide RNAs and primers used to generate the CRISPR edited animals.

**Table 4.**
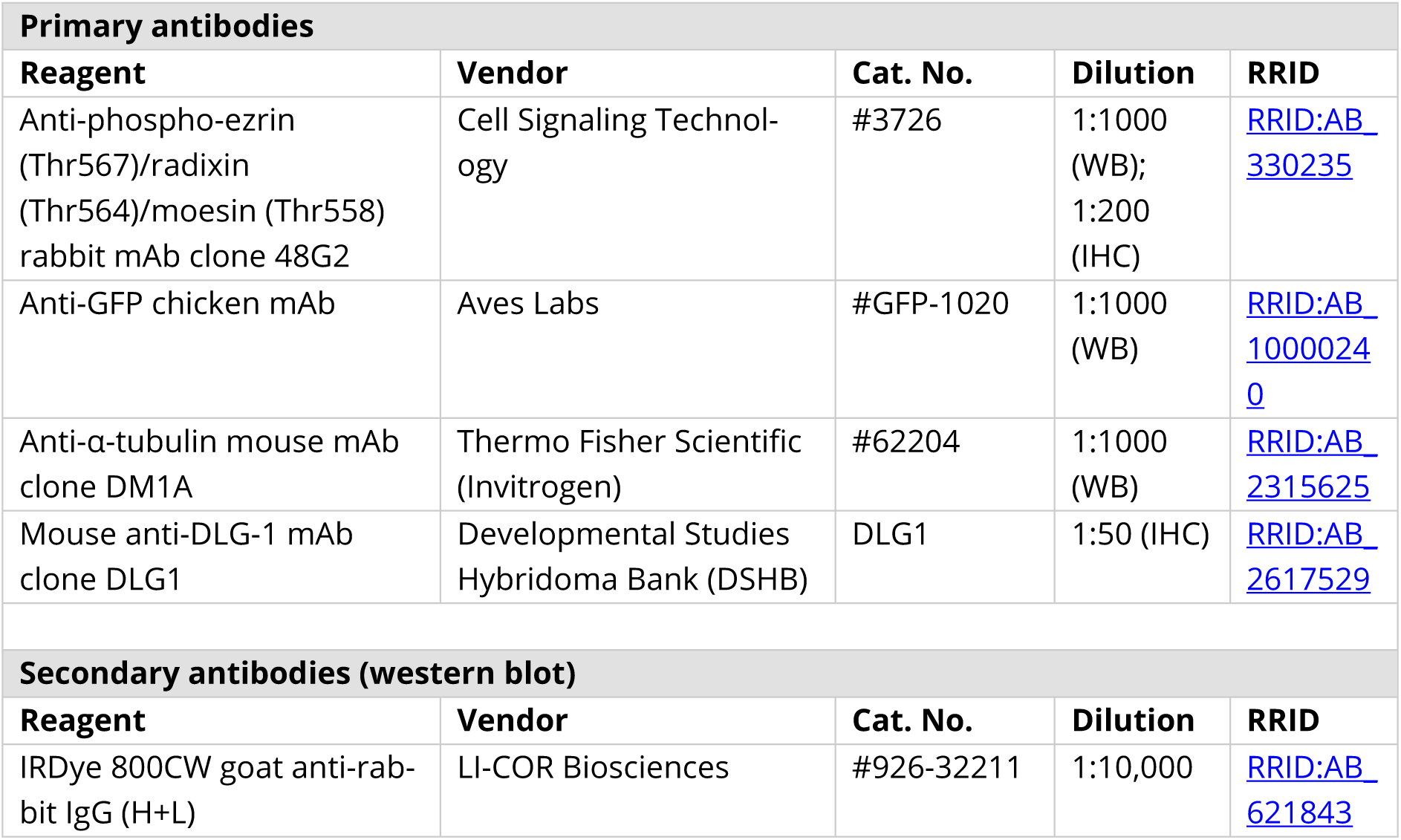
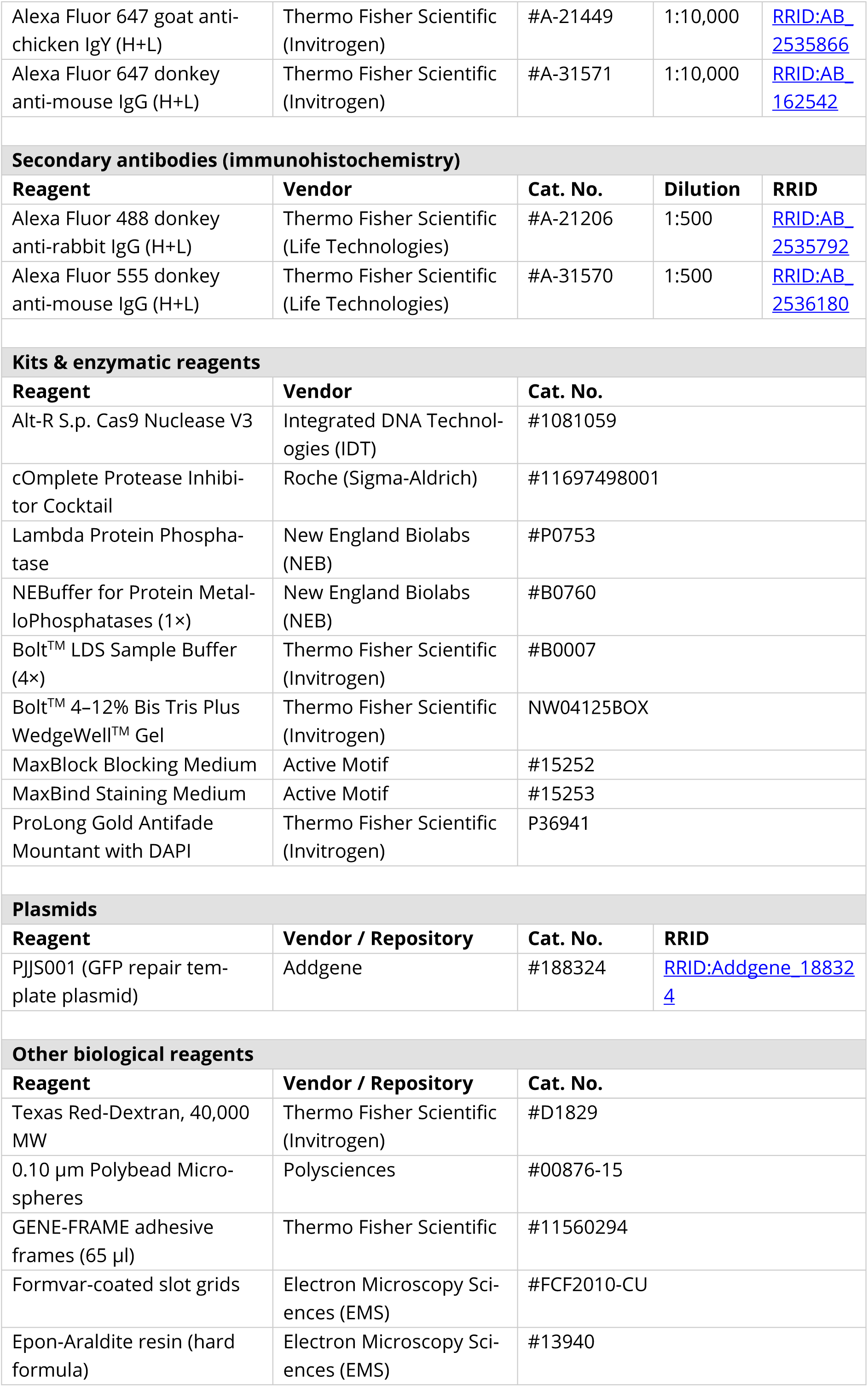
List of reagents used in this study.

### RNAi feeding

RNAi feeding was performed as previously described (Ramalho et al., 2020). L4 hermaphrodites were transferred to NGM-RNAi plates against target genes, and phenotypes were analyzed in the F1 generation. Table 1 contains a list of targeted genes including the corresponding RNAi library.

### Texas Red-Dextran feeding assay

L1 worms were collected and washed three times in M9 buffer (0.22 M KH_2_PO_4_, 0.42 M Na_2_HPO_4_, 0.85 M NaCl and 0.001 M MgSO_4_). The worms were then pelleted and transferred to a 1 mg/ml Texas Red-Dextran 40,000 MW (Thermo Fisher D1829) solution in egg buffer (118 mM NaCl, 48 mM KCl, 2 mM MgCl_2_, 2 mM CaCl_2_, 25 mM HEPES pH 7.3) and incubated for 3 hours at room temperature gently rocking on a tilting plate. The worms were then washed three times in M9 to remove the dye in solution. The animals were paralyzed in 20 mM tetramisole on an agarose pad on a glass slide and imaged using spinning disc microscopy.

### Western blot analysis

For the quantification of ERM-1 T544 phosphorylation, animals were grown on 6 cm NGM plates with OP50 bacteria. Adult animals were washed off full plates with M9 buffer supplemented with 0.05% Tween-20 (M9+T). The animals were washed an additional two times in M9+T and bleached two times in bleaching solution (0.25 M NaOH and ∼0.21 M sodium hypochlorite in distilled water) to harvest embryos. The bleaching solution was diluted with M9 medium and then the embryos were washed three times in M9. Embryos were incubated overnight at room temperature in M9 medium and filtered with a 10 µm filter to collect animals in the first larval stage. L1 larvae were washed in ice-cold Western blot (WB) lysis buffer containing 25 mM Tris-HCl pH 7.5, 150 mM NaCl, 1 mM EDTA, 0.5% IGEPAL CA-630 (Sigma-Aldrich), 1X cOmplete Protease Inhibitor Cocktail (Roche), and 50 mM NaF. Animals were then lysed in 100–200 µl WB lysis buffer followed by sonication for 10 min (sonication cycle: 30 s ON, 30 s OFF) at 4°C in a Bioruptor ultrasonication bath (Diagenode) at high energy setting. Lysates were cleared by centrifugation and denatured in 1x LDS Sample Buffer (Thermo Fisher) with 50 mM DTT at 99 °C for 10 min. The protein lysate was resolved on precast Bolt^TM^ 4–12% Bis Tris Plus WedgeWell^TM^ Gels (Invitrogen) before being transferred to a polyvinylidene difluoride membrane (pore size 0.45 µm) by wet transfer with a Mini Trans-Blot Cell (Bio-Rad). After transfer, blots were blocked for 1 hour at room temperature with blocking solution (5% BSA in TBST). Blots were probed overnight at 4°C in blocking solution with the following primary antibodies: anti-phospho-ezrin (Thr567)/radixin (Thr564)/moesin (Thr558) (48G2) rabbit mAb (#3726, Cell Signaling Technology; 1:1000), anti-GFP chicken mAb (#GFP-1020, Aves Labs; 1:1000), and anti-tubulin mouse mAb (#62204, Thermo Fisher; 1:1000). Blots were then washed three times with TBST for 10 min and incubated for 1 h at room temperature in the dark with the following secondary antibodies: anti-rabbit IRDye 800CW (#926-32211, LI-COR Biosciences; 1:10,000), Alexa Fluor 647 goat anti-chicken (#A21449, Thermo Fisher; 1:10,000), and Alexa Fluor 647 donkey anti-mouse (#A31571, Thermo Fisher; 1:10,000). Membranes were washed 3 times with TBST and kept in the dark before imaging on an Odyssey Infrared Imaging system (LI-COR Biosciences). The adjusted pERM density per sample was quantified using the Fiji 2.16.0/1.54p software. Background signal was subtracted using a 10px rolling ball background correction and the pERM density was corrected for loading differences using the ERM-1::GFP density. The adjusted values were normalized to the average control value.

### Phosphatase treatment

To test the anti-phospho-ezrin antibody, blocked Western blots were washed three times with TBST and treated with 2000 units Lambda Protein phosphatase (NEB) in phosphatase buffer (1X NEBuffer Pack for Protein MetalloPhosphatases supplemented with 1 mM MnCl_2_) for 60 min at 30°C in a humid chamber. As controls, blots were incubated in either TBST or phosphatase buffer. The blots were then washed three times with TBST and stained with primary and secondary antibodies according to the Western blot protocol.

### Immunofluorescence staining

For the staining of embryos, embryos were obtained by bleaching adult worms (see Western blot analysis for bleaching protocol) and let develop for 3 hours at room temperature in M9 medium. To get rid of salts in solution, the embryos were washed three times in MQ-water before being transferred to poly-L-lysine-coated frosted glass slides. A #1 coverslip (Carl Roth) was lowered on top of larvae/embryos, followed by freezing in liquid nitrogen and snapping off the coverslip. Fixation was performed in cold formaldehyde solution with phosphatase inhibitors: 1.6% formaldehyde (Sigma-Aldrich), 250 μM EDTA, and 50 mM NaF in PBS (1,35 M NaCl, 27 mM KCl, 100 mM Na_2_HPO_4_, 18 mM KH_2_PO_4_) at room temperature for 10 min. Samples were washed three times in wash buffer (250 μM EDTA, 50 mM NaF and 0.1% Triton X-100 from Sigma-Aldrich in PBS) and permeabilized in permeabilization buffer (0.5% Triton X-100, 250 µM EDTA and 50 mM NaF in PBS) for 30 min at room temperature. Samples were then washed one time in wash buffer and two times in PBS, and blocked in MaxBlock Blocking Medium (Active Motif, #102224) for 1 hour at room temperature in a humid chamber. The following primary antibodies were applied in MaxBind Staining Medium (Active Motif, #102227) overnight at 4°C in a humid chamber: anti-phospho-ezrin (Thr567)/radixin (Thr564)/moesin (Thr558) (48G2) rabbit mAb (#3726, Cell Signaling Technology, 1:200); and mouse anti-DLG-1 (DLG1, Hybridoma bank, RRID:AB_2617529, 1:50). Samples were then washed three times in PBS for 10 min and stained with the following secondary antibodies in MaxBind Staining Medium for 1 hour at room temperature in the dark: Alexa-Fluor 488 donkey anti-rabbit (A-21206, Life Technologies, 1:500) and Alexa-Fluor 555 donkey anti-mouse (A-31570, Life Technologies, 1:500). Samples were then washed three times in PBS for 10 min in the dark, mounted with Prolong Gold Antifade with DAPI (Thermo Fisher) under a coverslip, and sealed with nail polish before being imaged using spinning disk microscopy.

### Spinning disk microscopy

Imaging of living animals was performed by mounting the larvae and embryos on a 5% agarose pad in 20 mM tetramisole solution in M9 (0.22 M KH_2_PO4, 0.42 M Na_2_HPO_4_, 0.85 M NaCl, 0.001 M MgSO_4_). To collect freshly hatched L1 stage larvae for imaging, 10-20 gravid adults were drop-bleached on empty NGM plates the evening prior. Spinning disk confocal imaging was performed using a Nikon Ti-U microscope equipped with a Yokogawa CSU-X1 spinning disc using a Nikon CFI Plan Apo λ 100x/1.45 0.13mm WD OIL 71, and a Prime BSI Express Scientific CMOS camera (Teledyne Photometrics). Spinning disk images were acquired using MetaMorph Microscopy Automation & Image Analysis Software (version 7.10.5.476). For the imaging of large gaps in L1 larvae, a Nikon CFI Plan Apo λ 60x/1.40 0.13mm WD OIL 94 objective was used. All stacks along the z-axis were obtained at 0.25-μm intervals, and maximum intensity Z projections were done in Fiji 2.16.0/1.54p software. Image scales were calibrated using a micrometer slide. For display in Figures all level adjustments, false coloring, and image overlays were done in Adobe Photoshop 2025 (version 26.10.0). Image rotation, cropping, and panel assembly were done in Adobe Illustrator 2025 (29.8.1). All edits were done nondestructively using adjustment layers and clipping masks, and images were kept in their original capture bit depth until final export from Illustrator for publication.

### Survival assay

To assess survival levels, the total progeny of 4 P0 animals per plate was scored after 24 hours, when the P0s were removed. The unhatched progeny were scored 24 h after removal, and the number of F1 adults were scored at 96 h after removal. The number of unhatched progeny represents embryonic lethality, the number of adults was scored as ‘survival’, and the remainder scored as larval lethality.

### Electron microscopy

To collect freshly hatched L1 stage larvae for imaging, 10-20 gravid adults were drop-bleached on empty NGM plates the evening prior. L1 stage worms were fixed using high-pressure freezing (HPF) with the HPM Live µ system (CryoCapCell) and subsequently freeze-substituted in anhydrous acetone containing 1% OsO_4_, 0.5% glutaraldehyde, and 0.25% uranyl acetate for 60 hours in a freeze substitution (FS) system (AFS-2, Leica Microsystems). The larvae were embedded in an Epon-Araldite resin mix (EMS hard formula). To improve anteroposterior orientation and facilitate sectioning, flat-embedding was performed using adhesive frames (11560294 GENE-FRAME, 65 µl, Thermo Fisher Scientific). Ultrathin sections were cut on an ultramicrotome (UC7, Leica Microsystems) and collected on formvar-coated slot grids (FCF2010-CU, EMS). Each larva was sectioned in five different places, with at least a 7 µm gap between each grid to ensure observation of different cells. Each grid contained a minimum of 5-10 consecutive sections, each 70 nm thick. The TEM grids were examined using a JEM-1400 electron microscope (JEOL) operating at 120 kV, equipped with a Gatan Orius SC200 camera (Gatan) and controlled by the Digital Micrograph software. The length and density of microvilli were quantified using Fiji on TEM images from at least five grids per worm.

### Fluorescence recovery after photobleaching (FRAP)

To collect fresh L1 larvae, 10–20 gravid adults were partially dissolved in 50–100 µl of bleaching solution (0.5 M NaOH and ∼0.21 M sodium hypochlorite in distilled water) on an NGM plate without bacteriaI. After overnight incubation at room temperature, the hatched L1 larvae were collected from the plates, paralyzed in 20 mM tetramisole for 15 min and mounted on 10% agarose pads in 0.10 µm Polybead Microspheres (Polysciences, #00876-15). FRAP imaging was performed using a Nikon Ti-U microscope equipped with a Yokogawa CSU-X1 spinning disc using a 100×-1.4 NA objective, and a Prime BSI Express Scientific CMOS camera. FRAP images were acquired using MetaMorph Microscopy Automation & Image Analysis Software. Timelapse videos (20s intervals, 20 min) were recorded prior, during and after photobleaching, and each frame was analyzed in Fiji. The ERM-1::GFP intensity was measured in and outside the bleached region using the peak histogram intensity of a 20-pixel wide line perpendicular to the apical intestinal membrane. Background values were subtracted from the ERM-1::GFP signals using the average intensity of a region outside the worm. To correct for acquisition photobleaching, ERM-1::GFP intensity measurements in the bleached region were divided by the measured intensity of the region of the bleached region normalized to the corresponding intensity of the five pre-photobleached frames. The corrected ERM-1::GFP intensity of the first post-bleach frame (t=0) was subtracted from all corrected intensities as a zero baseline. The change in corrected intensity normalized against pre-photo-bleached intensities corresponded to fluorescence recovery. Two FRAP measurements were averaged per worm.

### Quantitative microscopy imaging analysis

Quantitative analysis of spinning disc images was done in Fiji 2.16.0/1.54p software. To determine the number of constrictions per worm, the number of points at the midline where no separation in IFB-2::mCherry and/or YFP::ACT-5 signal was observed was quantified. Larger lumen gaps were detected in PAR-6::GFP apical signal or DLG-1::mCherry apical junctional signal. The number of gaps are a sum of gaps observed within the intestine and between the intestine and pharynx. The localization of these large gaps was determined based on the expression of the *pGES-1::*PH::BFP; *pGES-1*::H2B::mCherry transgene in the BOX1408 strain.

Intensity distribution profiles to analyze co-distribution of GCK-4::GFP and ERM-1::mCherry::AID* in the microvilli were obtained by taking 15px-wide line scans perpendicular to the apical membrane spanning both lumen membranes. The line scans were split in half, the left part was mirrored to match the right half and all scans were aligned to the peak in GCK-4::GFP signal and min-max normalized. The width of the lumen and intestine was obtained using the *par*-*6::GFP*; *let*-*143::mCherry* strains. The width measurement was performed on maximum projections of the PAR-6::GFP. A single 15 pixel wide line was drawn along the midline and width was measured using the VasoMetrics macro(McDowell et al., 2021). A 100 pixel length was chosen for the perpendicular lines, with 1 µm distance between the crosslines.

To measure the junctional intensities of HMR-1::GFP and DLG-1::mCherry in 1.5-fold stage embryos, a maximum intensity projection of 5 slices was analyzed. A 15-pixel wide line perpendicular to the junctions was taken to obtain the max value along the line and the final value is the average of three max measurements per animal. The average value was corrected for imaging system background levels by subtracting the average of a region with the field of view that did not contain any animals or debris. The ratio of HMR-1 to DLG-1 intensity levels was calculated by dividing the background corrected averages.

To measure the apical YFP::ACT-5, ERM-1::GFP, GCK-4::GFP and LifeAct::mRuby levels during development, a maximum projection of 3 slices for bean, 1.5-fold stage embryos and L1 larvae, and a single slice view for 2-fold stage embryos were analyzed. A region outside of the animal was chosen for background subtraction and the average was subtracted from the image. Next, a line with 25 pixel width was drawn from the first intestinal ring to the 3^rd^–4^th^ ring and straightened using the Straighten function. The maximum value was determined for a line perpendicular to the lumen for each pixel along the X-axis. The maximum values were averaged and normalized to the average of control bean stage embryos, except for larvae where the data was normalized to the larval control.

### Phylogenetic tree reconstruction

For the collection of GCK-4 homologs a set of 15 Metazoan proteomes that span major metazoan lineages were downloaded from EukProt v3 (https://evocellbio.com/eukprot/) (Sobala, 2024). The species downloaded were *Amphimedon queenslandica, Caenorhabditis elegans, Clytia hemisphaerica, Drosophila melanogaster, Homo sapiens, Mnemiopsis leidyi, Mus musculus, Nematostella vectensis, Oscarella pearsei, Pleurobrachia bachei, Strongylocentrotus purpuratus, Sycon ciliatum, Tribolium castaneum, Trichoplax adhaerens, Hydra vulgaris*. We first built a hidden Markov model (HMM) of the kinase domain of GCK-4 homologs. To do that, a single Jackhmmer iteration (HMMER 3.4) (Potter et al., 2018) was performed with an e-value threshold of 1e^−65^ against all proteomes using the *C. elegans* GCK-4 as a query. The retrieved sequences were aligned using the MAFFT-L-INS-I v7.526 (Katoh and Standley, 2013) and the region corresponding to the kinase domain was extracted based on the Alphafold (Jumper et al., 2021) structure prediction of *C. elegans* GCK-4. The kinase domain region was then trimmed using trimAl v1.4.rev15 (-gt 0.6) to remove poorly aligned, gap-rich positions (Capella-Gutiérrez et al., 2009), and the resulting alignment was used to generate the HMM profile with HMMBUILD (HMMER v3.3.2) (Eddy, 2011). The HMM profile was then used to perform a HMMSEARCH (HMMER 3.4) with an e-value threshold of 1e^−60^ against all the proteomes. All the sequences were extracted and aligned with MAFFT-L-INS-I v7.526 and poorly aligned, gap-rich positions were removed by using trimAl (-gt 0.7) and subsequent manual visual inspection of the alignment. The phylogenetic tree was constructed with IQ-TREE v 2.3.6 (Minh et al., 2020) using the best fitted substitution model Q.insect+R7 (-m MFP) and ultrafast bootstrap with 1000 replicates (-B). The maximum likelihood tree was inferred as unrooted. Protein sequences linked to the same gene were detected and removed only for the GCK-4 OG. For visualization purposes, the tree was displayed with the root placed adjacent to the LOK/SLK orthologous group, and clades outside this group were collapsed for clarity. Tree was visualized using Interactive Tree of Life (iToL) (Letunic and Bork, 2021) and EukProt sequences were annotated using the NCBI identifiers. The final alignment FASTA file and IQ-TREE output in Newick format are available as supplementary files 2 and 3.

### Statistical analysis

All statistical analyses were performed using GraphPad Prism 10.4.1 or R. The sample sizes, statistical test used, and p values are indicated in the Figures and their legends. No statistical method was used to pre-determine sample sizes. No samples or animals were excluded from analysis. The experiments were not randomized, and the investigators were not blinded to allocation during experiments and outcome assessment. All presented graphs were made using GraphPad Prism and Adobe Illustrator. All tests were two-tailed.

For population comparisons, normality was first assessed using the D’Agostino & Pearson test. In all experiments, at least one population did not pass the normality test, hence data were treated as non-normally distributed and compared using a Mann-Whitney test. For comparisons of lumen enrichment gap counts between groups, Fisher’s exact test was used. Normalized Western blot intensity values were analyzed using a one-sample t-test, testing whether the mean differed from the theoretical value of 1 for each condition.

## Supporting information

File S1 - DNA sequence files

File S2 - FASTA alignment

File S3 - IQ-TREE Newick file

File S4 - Raw data of all quantified figures

## Supplementary material

This publication is accompanied by the following supplementary material:

- Supplementary Figure 1: Loss of GCK-4, the *C. elegans* ortholog of mammalian LOK/SLK kinases, leads to digestive tract blockages. Related to Figure 1.
- Supplementary Figure 2: GCK-4 is expressed in multiple epithelial tissues, and kinase-dead GCK-4::GFP localizes to the apical luminal domain. Related to Figure 2.
- Supplementary Figure 3: lumen narrowing in *gck-4(null)* comma stage embryos. Related to Figure 3.
- Supplementary Figure 4: Junctional levels of HMR-1 and DLG-1 and large midline gaps in *gck-4(null)* embryos. Related to Figure 4.
- Supplementary Figure 5: Large lumen gap between pharynx and intestine in *gck-4(null)* L1 larvae. Related to Figure 5.
- Supplementary Figure 6: Lower apical levels of ACT-5 and ERM-1 in *gck-4(null)* L1 larvae. Related to Figure 6.
- Supplementary Figure 7: α-pERM antibody phosphatase treatment on western blot membrane and survival assay including ERM-1(T544) phosphorylation mutants and mimetics. Related to Figure 7.
- Supplementary File 1: DNA sequence files of CRISPR alleles generated in this manuscript.
- Supplementary File 2: FASTA alignment file of GCK-4 homologs.
- Supplementary File 3: IQ-TREE Newick file of GCK-4 phylogenetic tree.
- Supplementary File 4: Raw data of all quantified figures.

## Data availability

All underlying numerical data for figures are provided in Supplementary File 4, including raw measurements, derived values, and the summary statistics plotted in the manuscript. All other data supporting the findings of this study are available within the paper and its supplementary information files. *C. elegans* strains are available upon reasonable request.

## Acknowledgements

We thank members of the M. Boxem, S. van den Heuvel, S. Ruijtenberg, S. Suijkerbuijk and L. Braccioli groups for helpful discussions. We thank G. Michaux and M. Galli for strains and reagents. Some strains were provided by the Caenorhabditis Genetics Center (CGC), which is funded by NIH Office of Research Infrastructure Programs (P40 OD010440). We also thank Wormbase (Sternberg et al., 2024) for providing accessible information concerning the genetics, genomics and biology of *C. elegans* and related nematodes. TEM Imaging was performed at the Microscopy Rennes Imaging Center (MRiC, Biosit, Rennes, France), a member of the National Infrastructure France-BioImaging supported by the French National Research Agency (ANR-10-INBS-04). Fluorescence microscopy imaging was performed at the Biology Imaging Center, Faculty of Sciences, Department of Biology, Utrecht University.

## Author contributions

- Mohamed Attaibi: Conceptualization, Formal analysis, Investigation, Visualization, Writing - Original Draft, Writing - Review & Editing.
- Jorian J. Sepers: Conceptualization, Formal analysis, Investigation, Visualization, Writing - Review & Editing.
- João J. Ramalho: Conceptualization, Formal analysis, Investigation, Visualization, Writing - Review & Editing.
- Ophélie Nicolle: Investigation, Visualization, Writing - Review & Editing.
- Lynn Hasenöhrl: Investigation, Visualization, Writing - Review & Editing.
- Savvas Tzavellas: Investigation, Visualization, Writing – Review & Editing.
- Ruben Schmidt: Investigation, Writing - Review & Editing.
- Grégoire Michaux: Supervision, Writing - Review & Editing, Funding acquisition.
- Mike Boxem: Conceptualization, Writing - Original Draft, Writing - Review & Editing, Supervision, Funding acquisition.

## References

Asan, A., Raiders, S. A. and Priess, J. R. Morphogenesis of the *C. elegans* Intestine Involves Axon Guidance Genes. PLoS Genet. 12, e1005950 (2016). 10.1371/journal.pgen.1005950

Baumgartner, M., Sillman, A. L., Blackwood, E. M., Srivastava, J., Madson, N., Schilling, J. W., Wright, J. H. and Barber, D. L. The Nck-interacting kinase phosphorylates ERM proteins for formation of lamellipodium by growth factors. Proc. Natl. Acad. Sci. U. S. A. 103, 13391–13396 (2006). 10.1073/pnas.0605950103

Bidaud-Meynard, A., Demouchy, F., Nicolle, O., Pacquelet, A., Suman, S. K., Plancke, C. N., Robin, F. B. and Michaux, G. High-resolution dynamic mapping of the *C. elegans* intestinal brush border. Dev. Camb. 148, (2021). 10.1242/dev.200029

Blasky, A. J., Mangan, A. and Prekeris, R. Polarized protein transport and lumen formation during epithelial tissue morphogenesis. Annu. Rev. Cell Dev. Biol. 31, 575–591 (2015). 10.1146/annurev-cellbio-100814-125323

Bovyn, M. and Haas, P. A. Shaping epithelial lumina under pressure. Biochem. Soc. Trans. 52, 331–342 (2024). 10.1042/BST20230632C

Brenner, S. The genetics of *Caenorhabditis elegans*. Genetics 77, 71–94 (1974).

Buckley, C. E. and St Johnston, D. Apical-basal polarity and the control of epithelial form and function. Nat. Rev. Mol. Cell Biol. 23, 559–577 (2022). 10.1038/s41580-022-00465-y

Capella-Gutiérrez, S., Silla-Martínez, J. M. and Gabaldón, T. trimAl: a tool for automated alignment trimming in large-scale phylogenetic analyses. Bioinformatics 25, 1972–1973 (2009). 10.1093/bioinformatics/btp348

Carberry, K., Wiesenfahrt, T., Windoffer, R., Bossinger, O. and Leube, R. E. Intermediate filaments in *Caenorhabditis elegans*. Cell Motil. Cytoskeleton 66, 852–864 (2009). 10.1002/cm.20372

Carberry, K., Wiesenfahrt, T., Geisler, F., Stöcker, S., Gerhardus, H., Überbach, D., Davis, W., Jorgensen, E., Leube, R. E. and Bossinger, O. The novel intestinal filament organizer IFO-1 contributes to epithelial integrity in concert with ERM-1 and DLG-1. Development 139, 1851–1862 (2012). 10.1242/dev.075788

Carreno, S., Kouranti, I., Glusman, E. S., Fuller, M. T., Echard, A. and Payre, F. Moesin and its activating kinase Slik are required for cortical stability and microtubule organization in mitotic cells. J. Cell Biol. 180, 739–746 (2008). 10.1083/jcb.200709161

Cavey, M. and Lecuit, T. Molecular bases of cell-cell junctions stability and dynamics. Cold Spring Harb. Perspect. Biol. 1, a002998 (2009). 10.1101/cshperspect.a002998

Chan, C. J. and Hiiragi, T. Integration of luminal pressure and signalling in tissue self-organization. Development 147, null (2020). 10.1242/dev.181297

Clarke, D. N. and Martin, A. C. Actin-based force generation and cell adhesion in tissue morphogenesis. Curr. Biol. CB 31, R667–R680 (2021). 10.1016/j.cub.2021.03.031

Coch, R. A. and Leube, R. E. Intermediate Filaments and Polarization in the Intestinal Epithelium. Cells 5, 32 (2016). 10.3390/cells5030032

Datta, A., Bryant, D. M. and Mostov, K. E. Molecular Regulation of Lumen Morphogenesis. Curr. Biol. 21, R126–R136 (2011). 10.1016/J.CUB.2010.12.003

Dickinson, D. J., Pani, A. M., Heppert, J. K., Higgins, C. D. and Goldstein, B. Streamlined Genome Engineering with a Self-Excising Drug Selection Cassette. Genetics 200, 1035–1049 (2015). 10.1534/genetics.115.178335

Dimov, I. and Maduro, M. F. The *C. elegans* intestine: organogenesis, digestion, and physiology. Cell Tissue Res. 377, 383–396 (2019). 10.1007/s00441-019-03036-4

Eddy, S. R. Accelerated Profile HMM Searches. PLOS Comput. Biol. 7, e1002195 (2011). 10.1371/journal.pcbi.1002195

Fehon, R. G., McClatchey, A. I. and Bretscher, A. Organizing the cell cortex: The role of ERM proteins. Nat. Rev. Mol. Cell Biol. 11, (2010). 10.1038/nrm2866

Feldman, J. L. and Priess, J. R. A role for the centrosome and PAR-3 in the hand-off of MTOC function during epithelial polarization. Curr. Biol. CB 22, 575–582 (2012). 10.1016/j.cub.2012.02.044

García-Alvarez, B., de Pereda, J. M., Calderwood, D. A., Ulmer, T. S., Critchley, D., Campbell, I. D., Ginsberg, M. H. and Liddington, R. C. Structural determinants of integrin recognition by talin. Mol. Cell 11, 49–58 (2003). 10.1016/s1097-2765(02)00823-7

Geisler, F., Coch, R. A., Richardson, C., Goldberg, M., Bevilacqua, C., Prevedel, R. and Leube, R. E. Intestinal intermediate filament polypeptides in C. elegans: Common and isotype-specific contributions to intestinal ultrastructure and function. Sci. Rep. 10, 3142 (2020). 10.1038/s41598-020-59791-w

Geisler, F., Remmelzwaal, S., Jankowski, V., Schmidt, R., Boxem, M. and Leube, R. E. Intermediate filament network perturbation in the C. elegans intestine causes systemic dysfunctions. eLife 12, e82333 (2023). 10.7554/eLife.82333

Ghaddar, A., Armingol, E., Huynh, C., Gevirtzman, L., Lewis, N. E., Waterston, R. and O’Rourke, E. J. Whole-body gene expression atlas of an adult metazoan. Sci. Adv. 9, eadg0506 (2023). 10.1126/sciadv.adg0506

Ghanta, K. S. and Mello, C. C. Melting dsDNA Donor Molecules Greatly Improves Precision Genome Editing in Caenorhabditis elegans. Genetics 216, 643–650 (2020). 10.1534/genetics.120.303564

Gloerich, M., ten Klooster, J. P., Vliem, M. J., Koorman, T., Zwartkruis, F. J., Clevers, H. and Bos, J. L. Rap2A links intestinal cell polarity to brush border formation. Nat. Cell Biol. 14, 793–801 (2012). 10.1038/ncb2537

Göbel, V., Barrett, P. L., Hall, D. H. and Fleming, J. T. Lumen Morphogenesis in *C. elegans* Requires the Membrane-Cytoskeleton Linker *erm-1*. Dev. Cell 6, 865–873 (2004). 10.1016/j.devcel.2004.05.018

Haas, M. A., Vickers, J. C. and Dickson, T. C. Rho kinase activates ezrin-radixin-moesin (ERM) proteins and mediates their function in cortical neuron growth, morphology and motility in vitro. J. Neurosci. Res. 85, 34–46 (2007). 10.1002/jnr.21102

Hall, D. H. and Altun, Z. F. Introduction. In WormAtlas, (2009). 10.3908/wormatlas.1.1

Hipfner, D. R., Keller, N. and Cohen, S. M. Slik Sterile-20 kinase regulates Moesin activity to promote epithelial integrity during tissue growth. Genes Dev. 18, 2243–2248 (2004). 10.1101/gad.303304

Hirokawa, N., Tilney, L. G., Fujiwara, K. and Heuser, J. E. Organization of actin, myosin, and intermediate filaments in the brush border of intestinal epithelial cells. J. Cell Biol. 94, 425–443 (1982). 10.1083/jcb.94.2.425

Hughes, S. C. and Fehon, R. G. Phosphorylation and activity of the tumor suppressor Merlin and the ERM protein Moesin are coordinately regulated by the Slik kinase. J. Cell Biol. 175, 305–313 (2006). 10.1083/jcb.200608009

Jewett, C. E. and Prekeris, R. Insane in the apical membrane: Trafficking events mediating apicobasal epithelial polarity during tube morphogenesis. Traffic (2018). 10.1111/tra.12579

Jumper, J., Evans, R., Pritzel, A., Green, T., Figurnov, M., Ronneberger, O., Tunyasuvunakool, K., Bates, R., Žídek, A., Potapenko, A., et al. Highly accurate protein structure prediction with AlphaFold. Nature 596, 583–589 (2021). 10.1038/s41586-021-03819-2

Katoh, K. and Standley, D. M. MAFFT Multiple Sequence Alignment Software Version 7: Improvements in Performance and Usability. Mol. Biol. Evol. 30, 772–780 (2013). 10.1093/molbev/mst010

Katzemich, A., Long, J. Y., Panneton, V., Fisher, L. A. B., Hipfner, D. and Schöck, F. Slik phosphorylation of Talin T152 is crucial for proper Talin recruitment and maintenance of muscle attachment in Drosophila. Development 146, dev176339 (2019). 10.1242/dev.176339

Kohler, P. O., Bridson, W. E., Hammond, J. M., Weintraub, B., Kirschner, M. A. and Van Thiel, D. H. Clonal lines of human choriocarcinoma cells in culture. Acta Endocrinol. Suppl. (Copenh.) 153, 137–153 (1971). 10.1530/acta.0.068s137

Kondrychyn, I., Kelly, D. J., Carretero, N. T., Nomori, A., Kato, K., Chong, J., Nakajima, H., Okuda, S., Mochizuki, N. and Phng, L.-K. Marcksl1 modulates endothelial cell mechanoresponse to haemodynamic forces to control blood vessel shape and size. Nat. Commun. 11, 5476 (2020). 10.1038/s41467-020-19308-5

Kunda, P., Pelling, A. E., Liu, T. and Baum, B. Moesin controls cortical rigidity, cell rounding, and spindle morphogenesis during mitosis. Curr. Biol. CB 18, 91–101 (2008). 10.1016/j.cub.2007.12.051

Letunic, I. and Bork, P. Interactive Tree Of Life (iTOL) v5: an online tool for phylogenetic tree display and annotation. Nucleic Acids Res. 49, W293–W296 (2021). 10.1093/nar/gkab301

Leung, B., Hermann, G. J. and Priess, J. R. Organogenesis of the Caenorhabditis elegans intestine. Dev. Biol. 216, 114–134 (1999). 10.1006/dbio.1999.9471

Lombardo, A. T., Mitchell, C. A. R., Zaman, R., McDermitt, D. J. and Bretscher, A. ARHGAP18-ezrin functions as an autoregulatory module for RhoA in the assembly of distinct actin-based structures. eLife 13, e83526 (2024). 10.7554/eLife.83526

Lubarsky, B. and Krasnow, M. A. Tube morphogenesis: making and shaping biological tubes. Cell 112, 19–28 (2003). 10.1016/s0092-8674(02)01283-7

Martin-Belmonte, F., Gassama, A., Datta, A., Yu, W., Rescher, U., Gerke, V. and Mostov, K. PTEN-mediated apical segregation of phosphoinositides controls epithelial morphogenesis through Cdc42. Cell 128, 383–397 (2007). 10.1016/j.cell.2006.11.051

Massarwa, R., Schejter, E. D. and Shilo, B. Z. Apical Secretion in Epithelial Tubes of the Drosophila Embryo Is Directed by the Formin-Family Protein Diaphanous. Dev. Cell 16, 877–888 (2009). 10.1016/J.DEVCEL.2009.04.010

Matsui, T., Maeda, M., Doi, Y., Yonemura, S., Amano, M., Kaibuchi, K., Tsukita, S. and Tsukita, S. Rho-kinase phosphorylates COOH-terminal threonines of ezrin/radixin/moesin (ERM) proteins and regulates their head-to-tail association. J. Cell Biol. 140, 647–657 (1998).

Matter, K. and Balda, M. S. SnapShot: Epithelial tight junctions. Cell 157, 992–992.e1 (2014). 10.1016/j.cell.2014.04.027

McDowell, K. P., Berthiaume, A. A., Tieu, T., Hartmann, D. A. and Shih, A. Y. VasoMetrics: unbiased spatiotemporal analysis of microvascular diameter in multi-photon imaging applications. Quant. Imaging Med. Surg. 11, 969–969 (2021). 10.21037/QIMS-20-920

Mège, R.-M., Gavard, J. and Lambert, M. Regulation of cell-cell junctions by the cytoskeleton. Curr. Opin. Cell Biol. 18, 541–548 (2006). 10.1016/j.ceb.2006.08.004

Minh, B. Q., Schmidt, H. A., Chernomor, O., Schrempf, D., Woodhams, M. D., von Haeseler, A. and Lanfear, R. Corrigendum to: IQ-TREE 2: New Models and Efficient Methods for Phylogenetic Inference in the Genomic Era. Mol. Biol. Evol. 37, 2461 (2020). 10.1093/molbev/msaa131

Mukenhirn, M., Wang, C.-H., Guyomar, T., Bovyn, M. J., Staddon, M., Veen, R. E. van der, Maraspini, R., Lu, L., Martin-Lemaitre, C., Sano, M., et al. Tight junctions control lumen morphology via hydrostatic pressure and junctional tension. Dev. Cell null, null (2024). 10.1016/j.devcel.2024.07.016

Naturale, V. F., Pickett, M. A. and Feldman, J. L. Persistent cell contacts enable E-cadherin/HMR-1- and PAR-3-based symmetry breaking within a developing C. elegans epithelium. Dev. Cell 58, 1830–1846.e12 (2023). 10.1016/j.devcel.2023.07.008

Navis, A. and Bagnat, M. Developing pressures: fluid forces driving morphogenesis. Curr. Opin. Genet. Dev. 32, 24–30 (2015). 10.1016/j.gde.2015.01.010

Ng, T., Parsons, M., Hughes, W. E., Monypenny, J., Zicha, D., Gautreau, A., Arpin, M., Gschmeissner, S., Verveer, P. J., Bastiaens, P. I., et al. Ezrin is a downstream effector of trafficking PKC-integrin complexes involved in the control of cell motility. EMBO J. 20, 2723–2741 (2001). 10.1093/emboj/20.11.2723

Pan, Y.-R., Tseng, W.-S., Chang, P.-W. and Chen, H.-C. Phosphorylation of moesin by Jun N-terminal kinase is important for podosome rosette formation in Src-transformed fibroblasts. J. Cell Sci. 126, 5670–5680 (2013). 10.1242/jcs.134361

Parisiadou, L., Xie, C., Cho, H. J., Lin, X., Gu, X.-L., Long, C.-X., Lobbestael, E., Baekelandt, V., Taymans, J.-M., Sun, L., et al. Phosphorylation of ezrin/radixin/moesin proteins by LRRK2 promotes the rearrangement of actin cytoskeleton in neuronal morphogenesis. J. Neurosci. Off. J. Soc. Neurosci. 29, 13971–13980 (2009). 10.1523/JNEUROSCI.3799-09.2009

Pelaseyed, T., Viswanatha, R., Sauvanet, C., Filter, J. J., Goldberg, M. L. and Bretscher, A. Ezrin activation by LOK phosphorylation involves a PIP2-dependent wedge mechanism. eLife 6, (2017). 10.7554/eLife.22759

Pickett, M. A., Sallee, M. D., Cote, L., Naturale, V. F., Akpinaroglu, D., Lee, J., Shen, K. and Feldman, J. L. Separable mechanisms drive local and global polarity establishment in the Caenorhabditis elegans intestinal epithelium. Dev. Camb. Engl. 149, dev200325 (2022). 10.1242/dev.200325

Pietromonaco, S. F., Simons, P. C., Altman, A. and Elias, L. Protein kinase C-theta phosphorylation of moesin in the actin-binding sequence. J. Biol. Chem. 273, 7594–7603 (1998). 10.1074/jbc.273.13.7594

Potter, S. C., Luciani, A., Eddy, S. R., Park, Y., Lopez, R. and Finn, R. D. HMMER web server: 2018 update. Nucleic Acids Res. 46, W200–W204 (2018). 10.1093/nar/gky448

Quizi, J. L., Baron, K., Al-Zahrani, K. N., O’Reilly, P., Sriram, R. K., Conway, J., Laurin, A.-A. and Sabourin, L. A. SLK-mediated phosphorylation of paxillin is required for focal adhesion turnover and cell migration. Oncogene 32, 4656–4663 (2013). 10.1038/onc.2012.488

Ramalho, J. J., Sepers, J. J., Nicolle, O., Schmidt, R., Cravo, J., Michaux, G. and Boxem, M. C-terminal phosphorylation modulates ERM-1 localization and dynamics to control cortical actin organization and support lumen formation during *Caenorhabditis elegans* development. Development 147, dev188011 (2020). 10.1242/dev.188011

Rambaud, B., Joseph, M., Tsai, F. C., De Jamblinne, C., Strakhova, R., Del Guidice, E., Sabelli, R., Smith, M. J., Bassereau, P., Hipfner, D. R., et al. Slik sculpts the plasma membrane into cytonemes to control cell-cell communication. EMBO J. 44, 2186–2186 (2025). 10.1038/S44318-025-00401-8

Remmelzwaal, S., Geisler, F., Stucchi, R., van der Horst, S., Pasolli, M., Kroll, J. R., Jarosinska, O. D., Akhmanova, A., Richardson, C. A., Altelaar, M., et al. BBLN-1 is essential for intermediate filament organization and apical membrane morphology. Curr. Biol. 31, 2334–2346.e9 (2021). 10.1016/j.cub.2021.03.069

Roch, F., Polesello, C., Roubinet, C., Martin, M., Roy, C., Valenti, P., Carreno, S., Mangeat, P. and Payre, F. Differential roles of PtdIns(4,5)P2 and phosphorylation in moesin activation during Drosophila development. J. Cell Sci. 123, 2058–2067 (2010). 10.1242/jcs.064550

Sadeghian, F., Ibrahim, I., Ravichandran, L., Henderson, G., Acharya, A., Wang, L., Lee, M. and Cram, E. J. An integrin binding motif in TLN-1/talin plays a minor role in motility and ovulation. MicroPublication Biol. 2023, (2023). 10.17912/micropub.biology.000726

Sallee, M. D., Pickett, M. A. and Feldman, J. L. Apical PAR complex proteins protect against programmed epithelial assaults to create a continuous and functional intestinal lumen. eLife 10, e64437 (2021). 10.7554/eLife.64437

Sepers, J. J., Ramalho, J. J., Kroll, J. R., Schmidt, R. and Boxem, M. ERM-1 Phosphorylation and NRFL-1 Redundantly Control Lumen Formation in the C. elegans Intestine. Front. Cell Dev. Biol. 10, 769862 (2022). 10.3389/fcell.2022.769862

Shaye, D. D. and Soto, M. C. Epithelial morphogenesis, tubulogenesis and forces in organogenesis. Curr. Top. Dev. Biol. 144, 161–214 (2021). 10.1016/bs.ctdb.2020.12.012

Shin, K., Fogg, V. C. and Margolis, B. Tight junctions and cell polarity. Annu. Rev. Cell Dev. Biol. 22, 207–235 (2006). 10.1146/annurev.cellbio.22.010305.104219

Sigurbjörnsdóttir, S., Mathew, R. and Leptin, M. Molecular mechanisms of de novo lumen formation. Nat. Rev. Mol. Cell Biol. 2014 1510 15, 665–676 (2014). 10.1038/nrm3871

Skop, A. R. and White, J. G. The dynactin complex is required for cleavage plane specification in early Caenorhabditis elegans embryos. Curr. Biol. CB 8, 1110–1116 (1998). 10.1016/s0960-9822(98)70465-8

Sobala, Ł. F. LukProt: A Database of Eukaryotic Predicted Proteins Designed for Investigations of Animal Origins. Genome Biol. Evol. 16, evae231 (2024). 10.1093/gbe/evae231

Sternberg, P. W., Van Auken, K., Wang, Q., Wright, A., Yook, K., Zarowiecki, M., Arnaboldi, V., Becerra, A., Brown, S., Cain, S., et al. WormBase 2024: status and transitioning to Alliance infrastructure. Genetics 227, iyae050 (2024). 10.1093/genetics/iyae050

ten Klooster, J. P., Jansen, M., Yuan, J., Oorschot, V., Begthel, H., Di Giacomo, V., Colland, F., de Koning, J., Maurice, M. M., Hornbeck, P., et al. Mst4 and Ezrin induce brush borders downstream of the Lkb1/Strad/Mo25 polarization complex. Dev. Cell 16, 551–562 (2009). 10.1016/j.devcel.2009.01.016

Tran Quang, C., Gautreau, A., Arpin, M. and Treisman, R. Ezrin function is required for ROCK-mediated fibroblast transformation by the Net and Dbl oncogenes. EMBO J. 19, 4565–4576 (2000). 10.1093/emboj/19.17.4565

Van Fürden, D., Johnson, K., Segbert, C. and Bossinger, O. The *C. elegans* ezrin-radixin-moesin protein ERM-1 is necessary for apical junction remodelling and tubulogenesis in the intestine. Dev. Biol. 272, 262–276 (2004). 10.1016/j.ydbio.2004.05.012

Viswanatha, R., Ohouo, P. Y., Smolka, M. B. and Bretscher, A. Local phosphocycling mediated by LOK/SLK restricts ezrin function to the apical aspect of epithelial cells. J. Cell Biol. 199, 969–984 (2012). 10.1083/jcb.201207047

Wald, F. A., Oriolo, A. S., Mashukova, A., Fregien, N. L., Langshaw, A. H. and Salas, P. J. I. Atypical protein kinase C (iota) activates ezrin in the apical domain of intestinal epithelial cells. J. Cell Sci. 121, 644–654 (2008). 10.1242/jcs.016246

Warner, A., Qadota, H., Benian, G. M., Vogl, A. W. and Moerman, D. G. The Caenorhabditis elegans paxillin orthologue, PXL-1, is required for pharyngeal muscle contraction and for viability. Mol. Biol. Cell 22, 2551–2563 (2011). 10.1091/mbc.E10-12-0941

Yang, H.-S. and Hinds, P. W. Increased ezrin expression and activation by CDK5 coincident with acquisition of the senescent phenotype. Mol. Cell 11, 1163–1176 (2003).

Yonemura, S., Matsui, T., Tsukita, S. and Tsukita, S. Rho-dependent and -independent activation mechanisms of ezrin/radixin/moesin proteins: an essential role for polyphosphoinositides in vivo. J. Cell Sci. 115, 2569–2580 (2002).

Zaman, R., Lombardo, A., Sauvanet, C., Viswanatha, R., Awad, V., Bonomo, L. E.-R., McDermitt, D. and Bretscher, A. Effector-mediated ERM activation locally inhibits RhoA activity to shape the apical cell domain. J. Cell Biol. 220, e202007146 (2021). 10.1083/jcb.202007146

Zhang, N., Khan, L. A., Membreno, E., Jafari, G., Yan, S., Zhang, H. and Gobel, V. The C. elegans Intestine As a Model for Intercellular Lumen Morphogenesis and In Vivo Polarized Membrane Biogenesis at the Single-cell Level: Labeling by Antibody Staining, RNAi Loss-of-function Analysis and Imaging. J. Vis. Exp. JoVE 56100 (2017). 10.3791/56100

Zhapparova, O. N., Fokin, A. I., Vorobyeva, N. E., Bryantseva, S. A. and Nadezhdina, E. S. Ste20-like protein kinase SLK (LOSK) regulates microtubule organization by targeting dynactin to the centrosome. Mol. Biol. Cell 24, 3205–3214 (2013). 10.1091/mbc.E13-03-0137

