## Supplementary Figures for "GCK-4 regulates apical actin organization and lumen formation in the *C. elegans* intestine"

**Condensed title:** Regulation of lumen formation by GCK-4

### Supplementary Figures

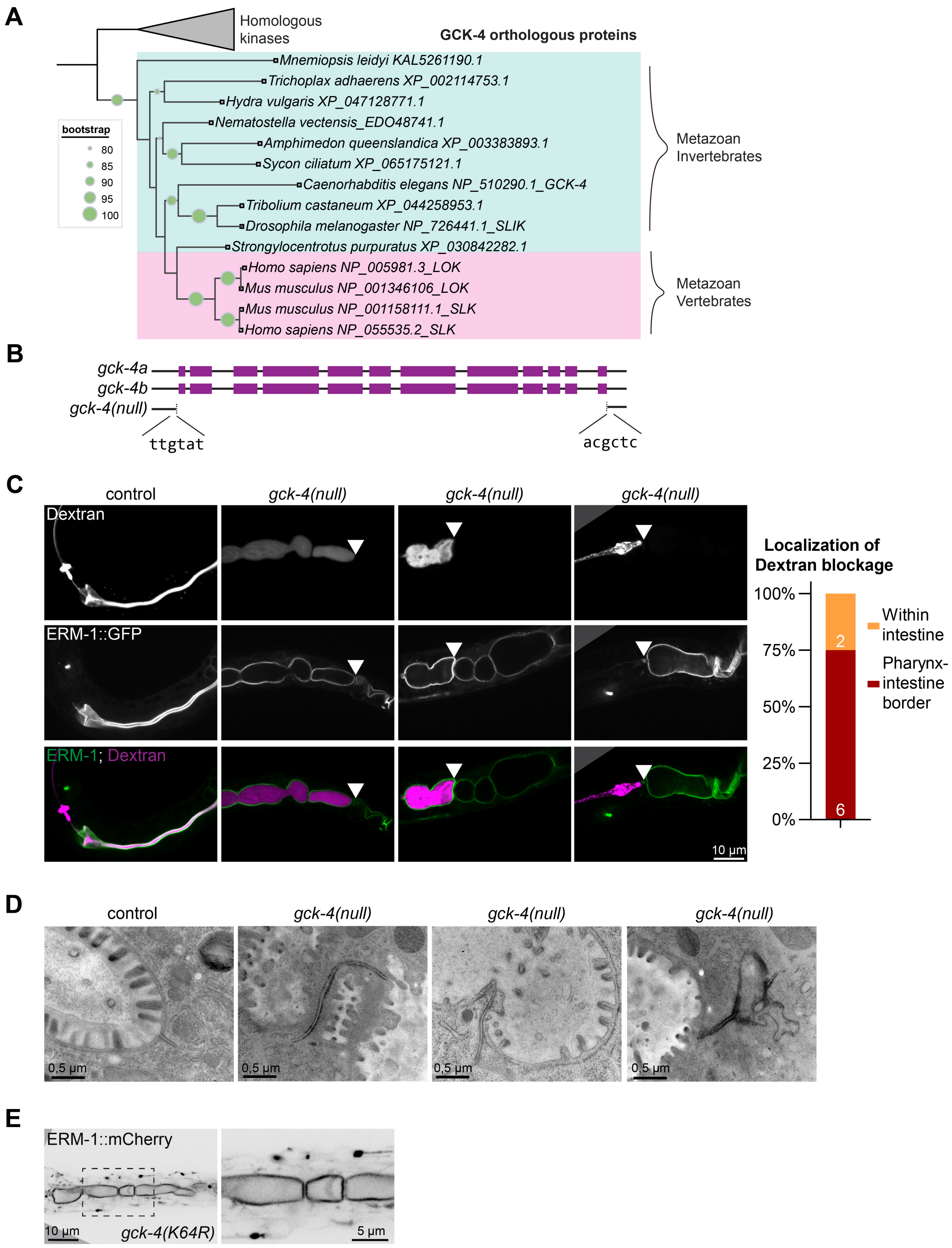


**Supplementary Figure 1: Loss of GCK-4, the *C. elegans* ortholog of mammalian LOK/SLK kinases, leads to digestive tract blockages.** (**A**) Phylogenetic tree showing the placement of GCK-4 within the LOK/SLK orthologous group (OG). The collapsed clade contains Metazoan OGs of 10 kinases homologous to GCK-4. Bootstrap analysis was performed using 1000 replicates, and support values ranging from 80 to 100 are indicated on the phylogenetic tree. The NCBI identifier is indicated in the label. (**B**) Schematic of *gck-4* isoforms and the deletion in the *gck-4(null)* allele. The isoforms only differ in the length of exon 10. (**C**) L1 larvae of the indicated genotypes carrying the apical marker ERM-1::GFP and fed with TexasRed-labeled Dextran. The white arrowheads indicate the constrictions that block the passage of dextran. All images are taken using a spinning-disk confocal microscope, and a single focal plane is shown. Quantifications of the localization of the Dextran blockages are shown in the graph next to the images. Pharynx-intestine border = Blockages observed posterior of the pharynx and anterior of the intestine, Within intestine = Blockages within the intestine. Strains used: BOX1030 and BOX1065. (**D**) Transmission Electron Microscope images of newly hatched *C. elegans* L1 larvae showing cell-cell junctions. Strains used: N2 and BOX1036. (**E**) Single plane internal view taken with a spinning disc confocal microscope showing the localization of ERM-1::mCherry::AID* in a *gck-4(K64R)* background. Enlarged version of the boxed region is shown to the right of the panel. Strain used: BOX1403.

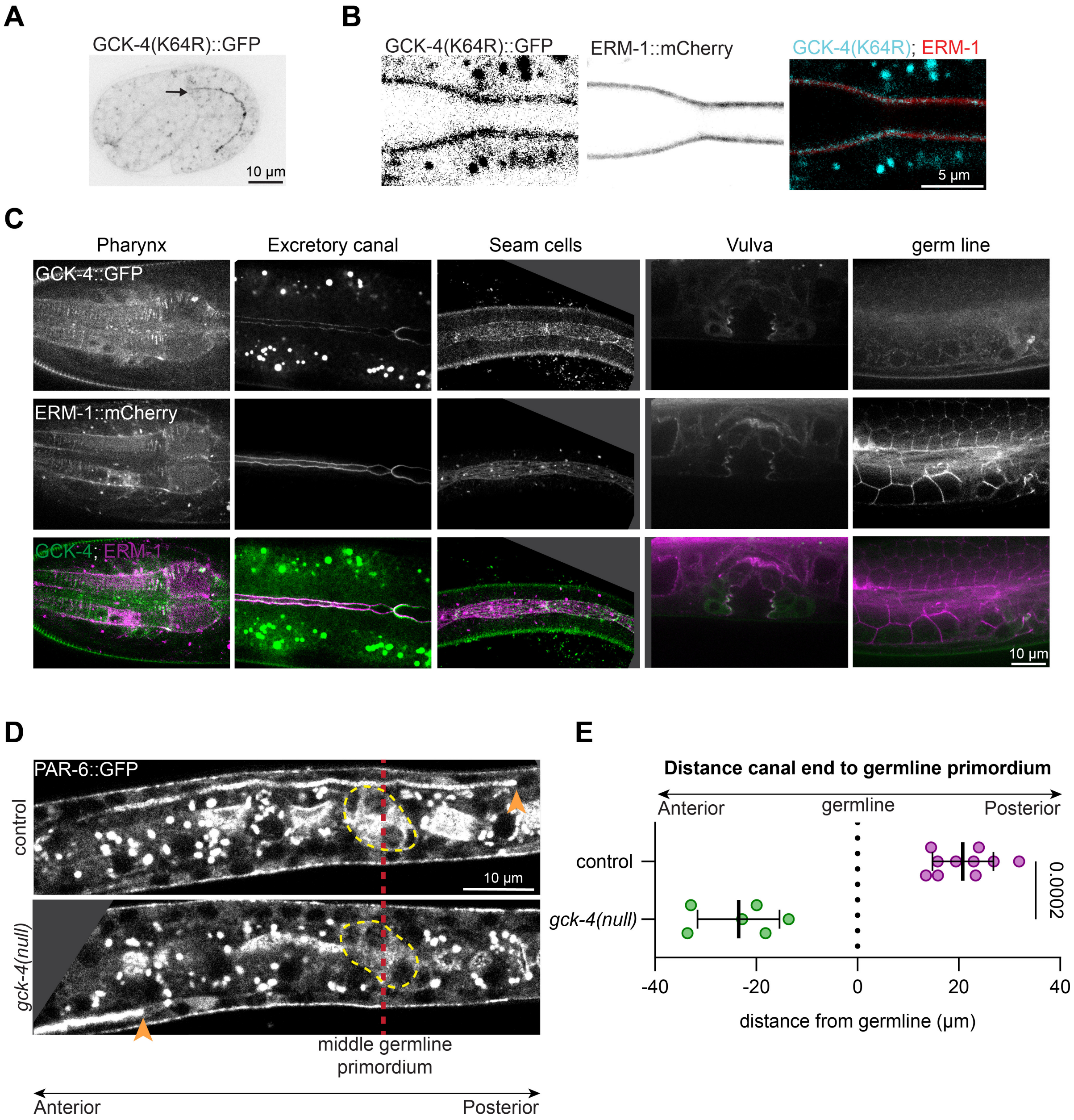


**Supplementary Figure 2: GCK-4 is expressed in multiple epithelial tissues, and kinase-dead GCK-4::GFP localizes to the apical luminal domain.** (**A**) Single focal plane at the center of the intestine showing the localization of GCK-4(K64R)::GFP in 1.6 fold embryo. Strain used: BOX1403. (**B**) Single focal plane at the center of the intestine showing the localization of GCK-4(K64R)::GFP in L1 larvae. Strain used: BOX1403. All images were taken using a spinning-disk confocal microscope. (**C**) Single focal plane showing localization of GCK-4::GFP and ERM-1::mCherry::AID* in indicated tissues. Strain used: BOX1095. (**D**) Single focal plane showing the localization of PAR-6::GFP in L1 larvae. Yellow outline shows the localization of the germline primordium, the red dashed line indicates the posterior end of the germline primordium, and the orange arrowheads the posterior end of the excretory canal. Strains used: BOX251 and BOX1129. (**E**) Quantifications of the distance from canal end to the middle of the germline primordium in larvae shown in Figure S2D. Total number of larvae analyzed in order of samples in graph: 6 and 10. Data are represented as mean ± SD and were analyzed with Mann-Whitney test; p value shown on graph. Strains used: BOX251 and BOX1129.

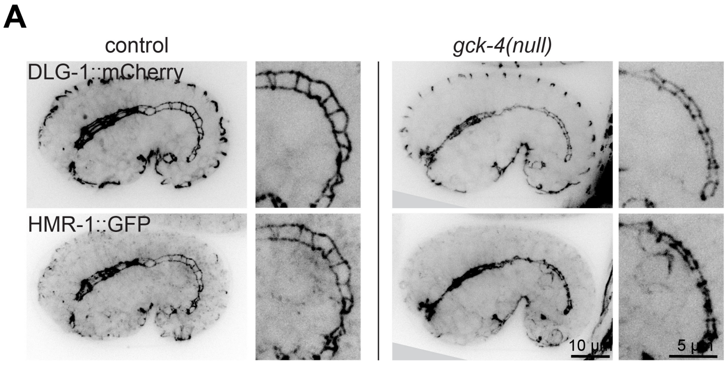


**Supplementary Figure 3: lumen narrowing in *gck-4(null)* comma stage embryos.** (**A**) Maximum intensity projections of 3D stacks taken with a spinning disc confocal microscope showing the localization of HMR-1::GFP and DLG-1::mCherry at the comma stage of embryonic development in control and *gck-4(null)* embryos. Enlarged versions of the boxed regions are shown to the right of each panel. Strains used: BOX1034 and BOX1067.

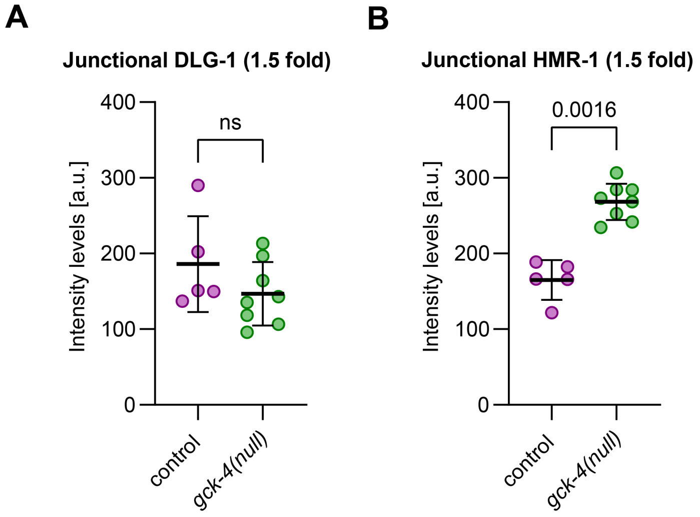


**Supplementary Figure 4: Junctional levels of HMR-1 and DLG-1 and large midline gaps in *gck-4(null)* embryos.** (**A, B**) Quantifications of junctional DLG-1::mCherry and HMR-1::GFP levels in 1.5-fold stage embryos as shown in (C). Data are represented as mean ± SD and were analyzed with Mann-Whitney test; p value shown on graph. Total number of embryos analyzed in order of samples in graphs: 5 and 8. Strains used: BOX1034 and BOX1067.

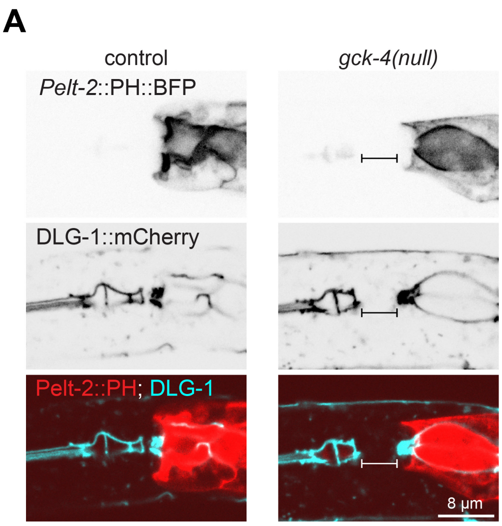


**Supplementary Figure 5: Large lumen gap between pharynx and intestine in *gck-4(null)* L1 larvae.** (**A**) Single focal planes taken with a spinning disc confocal microscope showing the localization of *Pelt-2*::PH::mTagBFP and DLG-1::mCherry in L1 larvae of indicated genotype. Bracket indicates large gap in pharyngeal–intestinal connection. Strains used: BOX1034 and BOX1067.

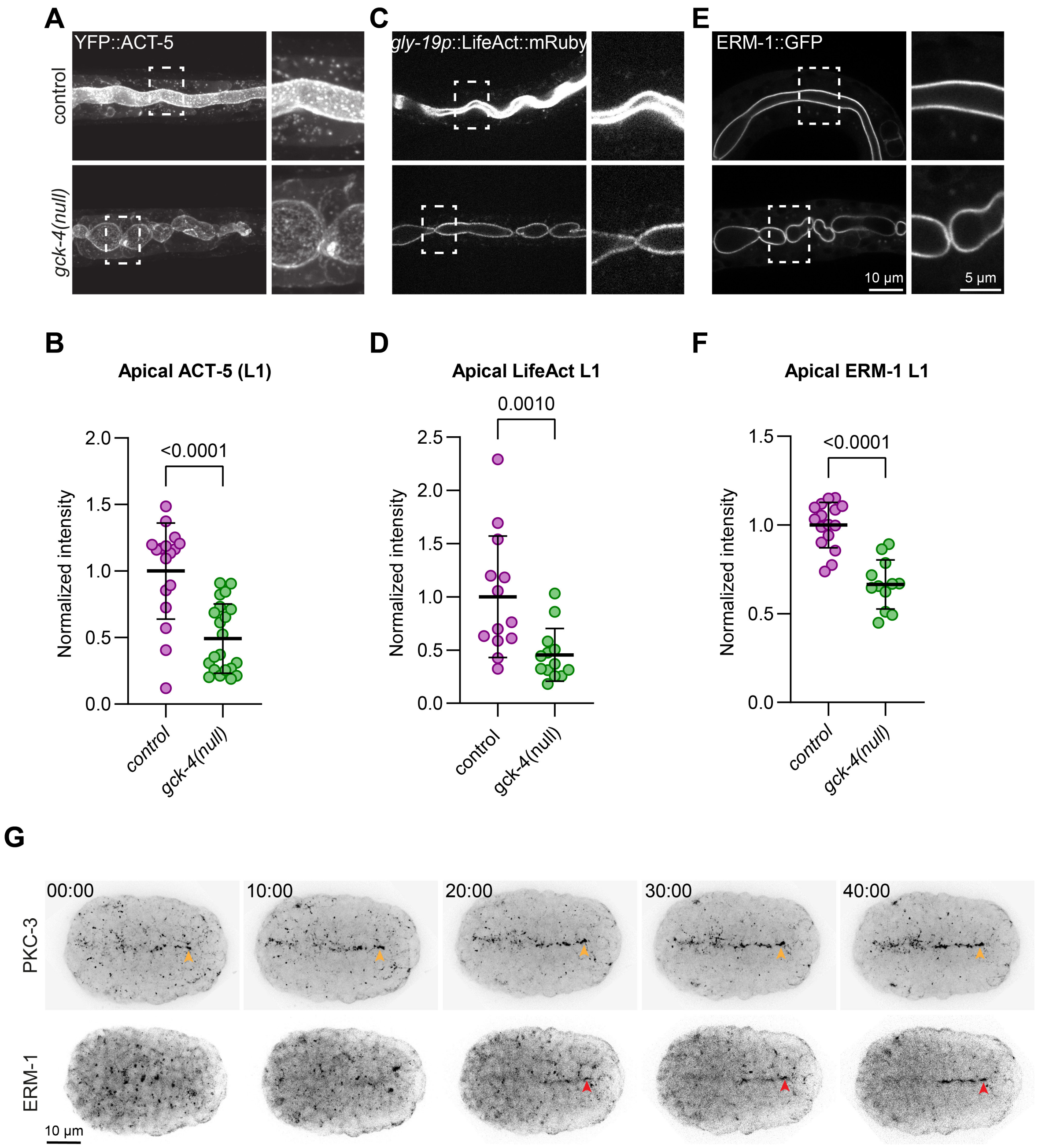


**Supplementary Figure 6: Lower apical levels of ACT-5 and ERM-1 in *gck-4(null)* L1 larvae.** (**A**) Maximum intensity projections of 3D stacks taken with a spinning disc confocal microscope showing the localization of *Pges-1*::YFP::ACT-5 in L1 larvae. Enlarged versions of the boxed regions are shown to the right of each panel. Strains used: BOX1031 and BOX1059. (**B**) Quantifications of apical YFP::ACT-5 levels in L1 larvae as shown in (A). Data are represented as mean ± SD and analyzed with Mann-Whitney test; p value shown on graph. Total number of larvae analyzed in order of samples in graphs: 17 and 21. (**C**) Single focal plane taken with a spinning disc confocal microscope showing the localization of *Pgly-19*::LifeAct::mRuby in L1 larvae. Enlarged versions of the boxed regions are shown to the right of each panel. Strains used: RHS43 and BOX1170. (**D**) Quantifications of apical *Pgly-19*::LifeAct::mRuby in L1 larvae as shown in Figure S6C. Data are represented as mean ± SD and analyzed with Mann-Whitney test; p value shown on graph. Total number of larvae analyzed in order of samples in graphs: 13 and 13. (**E**) Single focal plane taken with a spinning disc confocal microscope showing the localization of ERM-1::GFP in L1 larvae. Enlarged versions of the boxed regions are shown to the right of each panel. Strains used: BOX1030 and BOX1065. (**F**) Quantifications of apical ERM-1::GFP in L1 larvae as shown in Figure S6E. Data are represented as mean ± SD and analyzed with Mann-Whitney test; p value shown on graph. Total number of larvae analyzed in order of samples in graphs: 16 and 12. (**G**) Maximum intensity projections of 3D live-imaging timelapse stacks of the localization of PKC-3::GFP and ERM-

1::mCherry::AID* during LPC migration and midline spreading. Time between each frame is 10 min and t=00m:00s is when an LPC (indicated by PKC-3::GFP spots) arrives at the midline. Orange arrowheads show the LPC containing PKC-3::GFP and the red arrowheads indicate ERM-1::mCherry accumulation at the LPC. Strain used: BOX846.

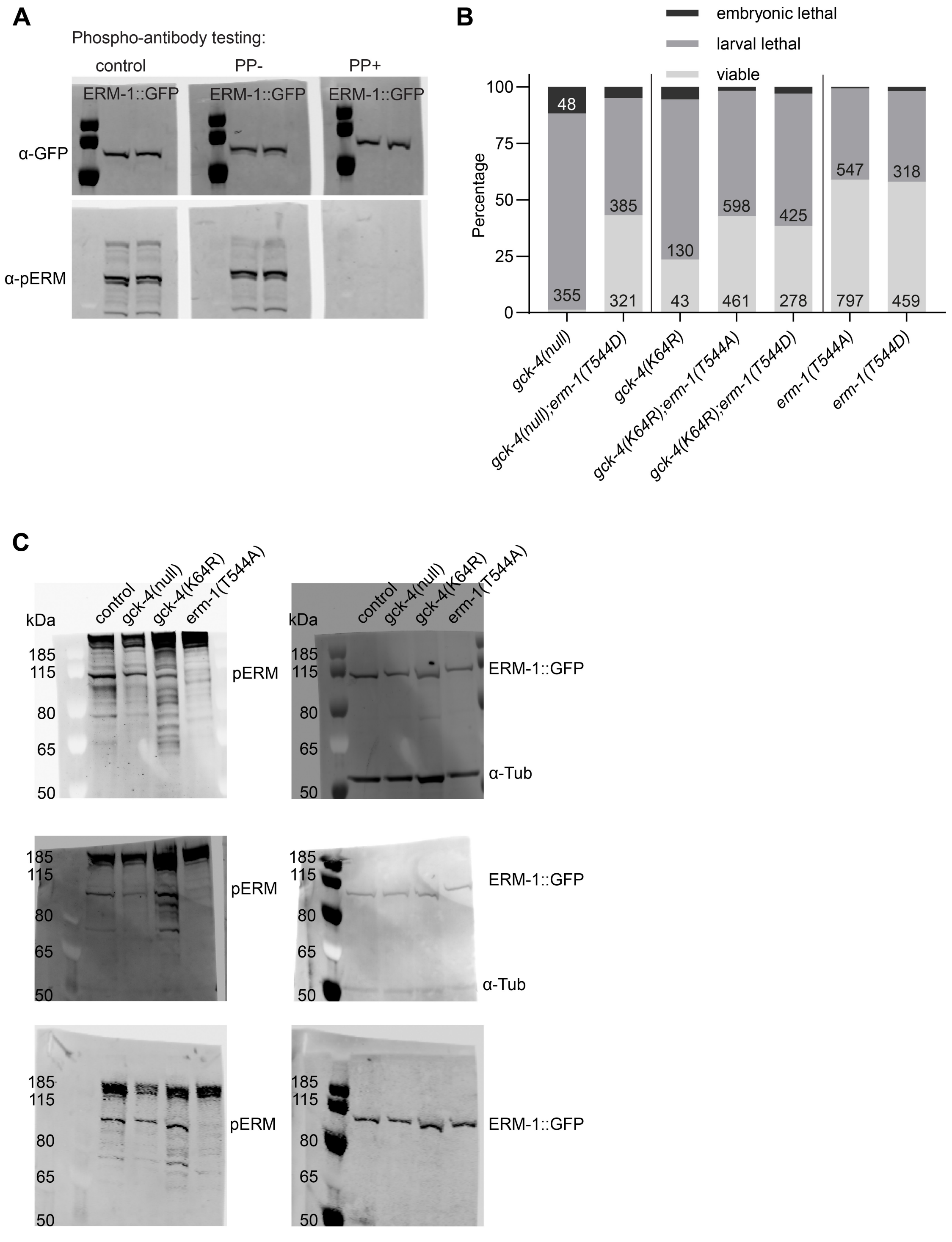


**Supplementary Figure 7: α-pERM antibody phosphatase treatment on western blot membrane and survival assay including ERM-1(T544) phosphorylation mutants and mimetics.** (**A**) Levels of ERM-1 T544 phosphorylation in freshly hatched L1 larvae assessed by western blot after lambda phosphatase treatment. Control = TBS, PP- = only PMP buffer, PP+ = PMP buffer + lambda phosphatase. α-pERM = phosphorylated ERM-1 T544, α-GFP = total ERM-1::GFP. Strain used: BOX1030. (**B**) Embryonic lethality, larval arrest, and survival to adulthood observed in progeny of animals of indicated genotypes. Data are represented as a proportion to the total amount of progeny, with the total number per outcome added to the graph. Total number of progeny counted in order of samples in graph: 408, 743, 183, 1078, 724, 1352, and 792. Strains used: BOX1036, BOX1252, BOX1402, BOX1404, BOX1405, BOX165 and BOX163. (**C**) Blots used for quantification of pERM levels in (A). Left column shows the pERM channel, right column the ERM-1::GFP channel. α-Tub = anti-tubulin antibody, α-pERM = phosphorylated ERM-1 T544, ERM-1::GFP = total ERM-1::GFP. Strains used: BOX1030, BOX1065, BOX1409 and BOX369.
